# Harnessing the Embryo Antitumor Factor NEPN for Broad-Spectrum Cancer Immunotherapy via Dual Modulation of Tumor Progression and Immune Suppression

**DOI:** 10.1101/2025.09.29.679137

**Authors:** Tzu-Pei Chang, Yu-Jui Ho, Mohamad Hamieh, Xin Huang, Nina Kim, Rajasekhar Vinagolu, Cheng Zhang, Rui Zhao, Shannon Mckinney-Freeman, Hu Li, Scott Lowe, Michel Sadelain, Kitai Kim

## Abstract

Modifying the tumor microenvironment to target malignant cells and spare healthy cells may be an alternative approach to cancer therapy. Early-stage embryos have been shown to reprogram metastatic tumor cells and prevent tumor formation of several types of cancers, including melanoma, leukemia, and neuroblastoma. However, embryonic factors that suppress tumors in early embryos have not been identified. Here we show that the embryonic secretory factor NEPN suppresses cancer progression and enhances tumor-infiltrating T cells by modulating HSP27 and ARG1 through TGFβ1 signaling, together with broader regulation of tumor-promoting and immune-modulating cytokines. Importantly, NEPN strengthens immune responses and cell death in not only non-solid tumor but also solid tumors by increasing T cell infiltration and activity in the tumor microenvironment. The ability of NEPN to boost immune responses within the tumor microenvironment establishes it as a promising therapeutic candidate for cancer immunotherapy.

## Main Text

During the early stages of embryogenesis, multiple proto-oncogenes, including c-myc, c-erb, c-ras, and c-src, are highly expressed to support rapid organ development^1^. The similar levels of these oncogenes expressed in adult somatic cells induce immortalization and oncogenic transformation; however, early-stage embryos rarely develop tumors, indicating the existence of a unique defense system to prevent tumor formation. A similar finding was observed in the human embryonic microenvironment by using human embryonic stem cells (hESC) as well^2^. Thus, we hypothesized that early embryos express one or more embryo-specific tumor suppressors to produce an embryonic microenvironment that inhibits cancer formation during embryo development.

In the 1980s, researchers injected melanoma, leukemia, or neuroblastoma cells into different developmental stages of mouse embryos and found that the early-stage embryonic tissue inhibits cancer development and induces tumor differentiation; but later-stage embryos or neonatal mice developed cancers.^3–7^ The mechanism remained unexplored due to the lack of genome-wide tools available at that time, we, therefore, hypothesized that the early embryo tissues might generate a specific microenvironment that contains unique secretory factors, capable of preventing cell linage-related tumor formation and inducing tumor differentiation of injected cancer cells. In addition, the embryonic fields might regulate tumors of their closely related cancers, in a tissue-specific manner, such as embryonic skin cells regulate melanoma^4^ or early hematopoietic tissues reprogram leukemic cells.^3^

### Early embryonic secretory protein NEPN functions as a tumor-suppressing factor

To investigate this unique system in the early embryonic microenvironment, we digested and cultured the dorsal embryonic skin cells from embryos at stage E9.5, E10.5, E12.5, and E14.5, the same sites where researchers had previously injected cancer cell lines.^3–7^ The resulting conditioned media (CM), which mimic the embryonic microenvironment, were collected and used to treat mouse B16 melanoma cells (**Fig.1a**). Treatment with E9.5 CM significantly reduced both the colony formation frequency and the number of cells per colony in mouse B16 melanoma cell (**Fig. 1b**) and increased cellular apoptosis (**Fig. 1c**). Interestingly, B16 cells treated with E9.5 CM exhibited morphological changes, becoming larger, darker, and more flattened compared to untreated cells, consistent with cancer cell differentiation observed in the embryonic microenvironment in previous reports (**Fig. 1d**).^3–7^ These characteristics are typical shown in differentiated melanoma cells, suggesting that the early embryonic microenvironment can inhibit cancer growth and involved in the induction of terminal tissue differentiation.^3–7^

**Figure 1.**
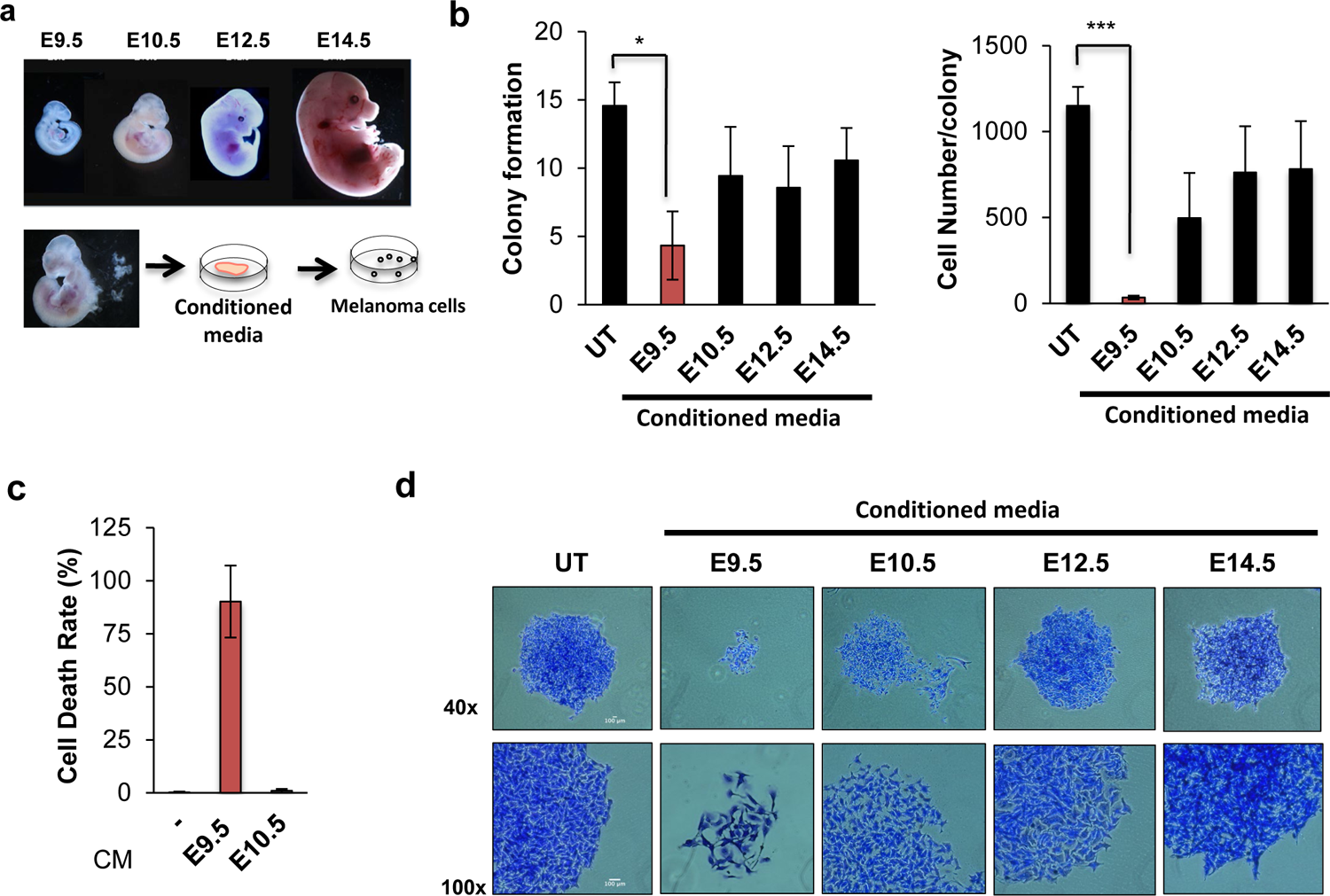
Identification of a unique system to inhibit cancer progression in the early embryonic environment. **a** Schematic of experiment set-up for collecting conditional medium (CM) at different embryonic stages. The conditioned medium was derived from embryonic dorsal skin tissues from different embryonic stages. Mouse B16-TGL melanoma cells were treated with this conditioned medium. Colony formation assay of B16-TGL melanoma cell was performed. **b,** Colony frequency (left) and the cell number of B16-TGL cells in each colony (right) were measured (n=5-6, biological using *in situ* cell death assay kit (n=3, biological independent experiments). **c,** Apoptosis in embryonic dorsal skin cells. Cell death rates were quantified using a TUNEL-based in situ cell death detection kit (Roche). Significant apoptosis was observed in cells treated with E9.5-conditioned media, whereas E10.5-conditioned media did not induce a detectable increase in cell death. A negative control (omission of terminal transferase enzyme) was included to confirm assay specificity. Data are presented as mean ± s.e.m. (n = 3, biologically independent experiments). **d,** the cellular morphology of B16-TGL cells was stained by crystal violet dye. * P < 0.05, * * P < 0.01, * * * P < 0.001,* * * * P < 0.0001(t-test). Scale bars, 100 μm.

To identify potential tumor-suppressing factors within the early embryonic microenvironment, we performed RNA-seq and qPCR analysis on dorsal embryonic skin cells from various embryonic stages. This analysis revealed NEPN, a secreted proteoglycan, as being highly expressed specifically in E9.5 embryonic skin cells (**Fig. 2a-b**). NEPN is a member of the small leucine-rich proteoglycan (SLRP) family,^8^ though its precise biological function remains largely uncharacterized. The limited published literature predicts that proteoglycan (such as Decorin), which are key functional molecules on the cell surface and in the pericellular microenvironment,^9–11^ might sequester excessive growth ligands such as TGFβ to modulate growth signaling pathways in the tumor microenvironment.^12^ HABP2 links NEPN to hyaluronan,^13^ and has recently been shown to exhibit anti-tumor activity by contributing to the formation of a specialized extracellular matrix in the cancer-resistant naked mole rat model (**Fig. 2c**),^14^ and was also identified as the second most highly expressed gene, specifically in E9.5 embryonic skin cells. (**Fig. 2a**).

**Figure 2.**
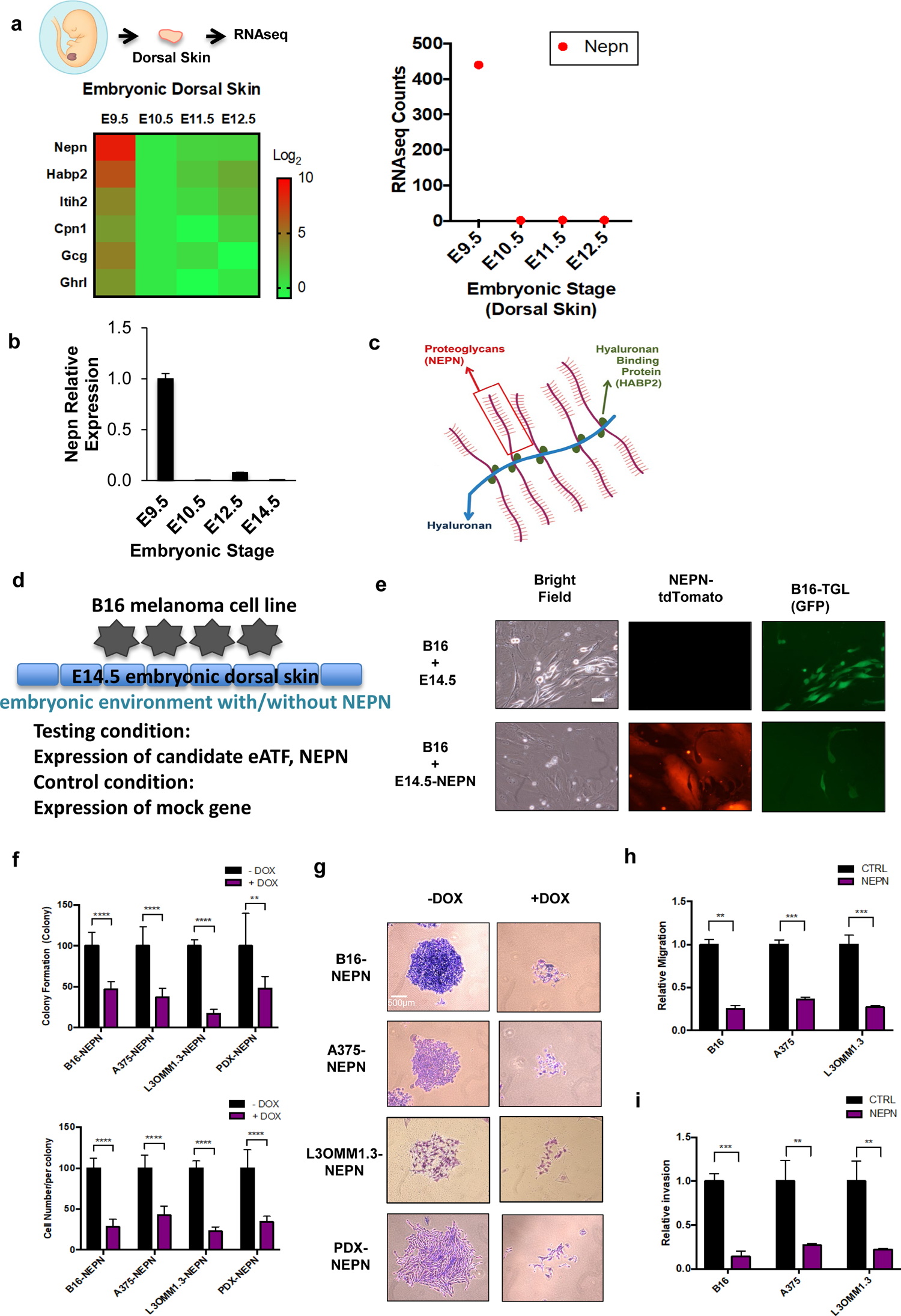
Identification of a secretory anti-tumor factor NEPN from the early embryonic stage. **a**, Heatmap (left) and count (right) of gene expression pattern (right) in dorsal skin cells at different embryonic stages by RNAseq analysis (n=2, independent experiments). **b,** RNA expression of NEPN in different embryonic-staged dorsal skin cells using qPCR (n=4, independent experiments). **c,** Schematic representation of NEPN as a proteoglycan in the extracellular matrix. NEPN (red) associates with the hyaluronan backbone (blue) via hyaluronan-binding proteins (HABP2, green), forming a structural signaling complex. **d,** Doxycycline-inducible co-culture system mimicking the early embryonic microenvironment. E14.5 dorsal skin cells were engineered with a doxycycline-inducible construct to express NEPN, recapitulating features of the E9.5 embryonic environment. The system allowed direct co-culture of embryonic skin cells with B16 melanoma cells to assess tumor-microenvironment interactions. **e,** Anti-tumor effect of NEPN in the co-culture system. E14.5 dorsal skin cells (tdTomato) were engineered to express NEPN and co-cultured with B16 melanoma cells (TGL, GFP). NEPN induction markedly suppressed melanoma growth while leaving normal embryonic skin cells unaffected, demonstrating cancer-specific activity. Scale bar, 100 μm. **f,** Frequency and cell number in each B16 colony were measured in different melanoma cell lines, mouse B16, human A375, human CDX L3OMM1.3, human PDX Mel1 cells overexpressed NEPN (n=6-8, independent experiments). **g,** Cellular morphology was stained by crystal violet dye. Scale bar, 500 μm. **h,** *In vitro* transwell invasion and **i,** migration assay of different melanoma cell lines expressing NEPN (n=3, independent experiments). * P < 0.05, * * P < 0.01, * * * P < 0.001,* * * * P < 0.0001(t-test).

Using a doxycycline-inducible co-culture model (**Fig. 2d**), E14.5 dorsal skin cells were engineered to express NEPN to mimic the E9.5 microenvironment. NEPN expression significantly inhibited B16 melanoma cell growth without detectable toxicity to normal skin cells. (**Fig. 2e**). These data confirm that NEPN is a key factor within the early embryonic microenvironment that regulates tumor progression by modulating the extracellular environment and the associated cellular signaling pathways.

We next assessed the direct effects of mouse NEPN on multiple melanoma models, including mouse B16 melanoma cells,^15^ human A375 melanoma cells,^16^ human patient cell-derived xenograft (CDX) melanoma L3OMM1.3 cells, and human patient-derived xenograft (PDX) melanoma Mel1 cells,^17^ using a lentivirus-doxycycline-inducible-NEPN system. NEPN overexpression significantly inhibited colony formation and cell proliferation across all tested melanoma cell lines (**Fig. 2f-g**) and reduced both the migration and invasion capabilities of melanoma cells across species (**Fig. 2h-i**).

To identify putative tumor-suppressing factors relevant to non-solid cancers such as leukemia, we analyzed the gene expression profile of the AGM (aorta-gonad-mesonephros) region, an early hematopoietic and leukemogenic niche associated with leukemia progression, across various embryonic stages using RNA-seq. Surprisingly, NEPN emerged among the most highly expression genes in the early-stage (E9.5) AGM region along with HABP2 (**Fig. S1a**). This finding suggests that NEPN may exert a conserved inhibitory effect on leukemia cells similar to its role in melanoma. Notably, E9.5 also marks the onset of T lymphoid lineage development,^18–21^ indicating that NEPN may play a role not only in T cell activation but also in the development of tumor-specific T cells. In support of this, our *in vitro* experiments demonstrated that NEPN inhibits the proliferation of NALM6 leukemia cells, while also promoting apoptosis and differentiation (**Fig. S1b-d**). Importantly, NEPN exhibited no cytotoxic effects on normal cells, including embryonic dorsal skin cells, melanocytes, and T cells (**Fig. S1e**).

### NEPN inhibits tumor proliferation and metastasis through the TGFβ1 signaling

NEPN is a secreted proteoglycan and a member of the SLRP family,^9–11^ known to bind and sequester growth ligands and receptors, thereby modulating growth signaling pathways within the tumor microenvironment.^12^ Yet, little is known about the function of NEPN. To define the mechanism of NEPN, we performed PTMScan-Mass Spectrometry analysis in A375 melanoma cells with and without NEPN expression. This analysis quantified 409 proteins and documented 1,006 unique phosphorylation and non-phosphorylation sites, allowing identification of the most significantly altered signaling pathways regulated by NEPN. Through this approach, we uncovered key post-translational modifications demonstrating how NEPN modulates tumor-suppressive pathways and immune-relevant signaling within the tumor microenvironment. The analysis revealed that Heat shock protein 27 (HSP 27, 85.7-fold change)^12,22,23^ and arginase 1 (ARG1, 72.1-fold change)^24–26^ were the most significantly altered targets (quantity) following NEPN expression (**Fig. 3a**). Notably, both HSP27 and ARG1 are regulated by a common upstream TGFβ signaling pathway.^26,27^ Based on these findings, we hypothesized that NEPN inhibits HSP27 and ARG1 through blocking ligands-receptor signals such as TGFβ by either sequestering ligands or blocking receptors) and consequently inhibits tumor formation and metastasis.

**Figure 3.**
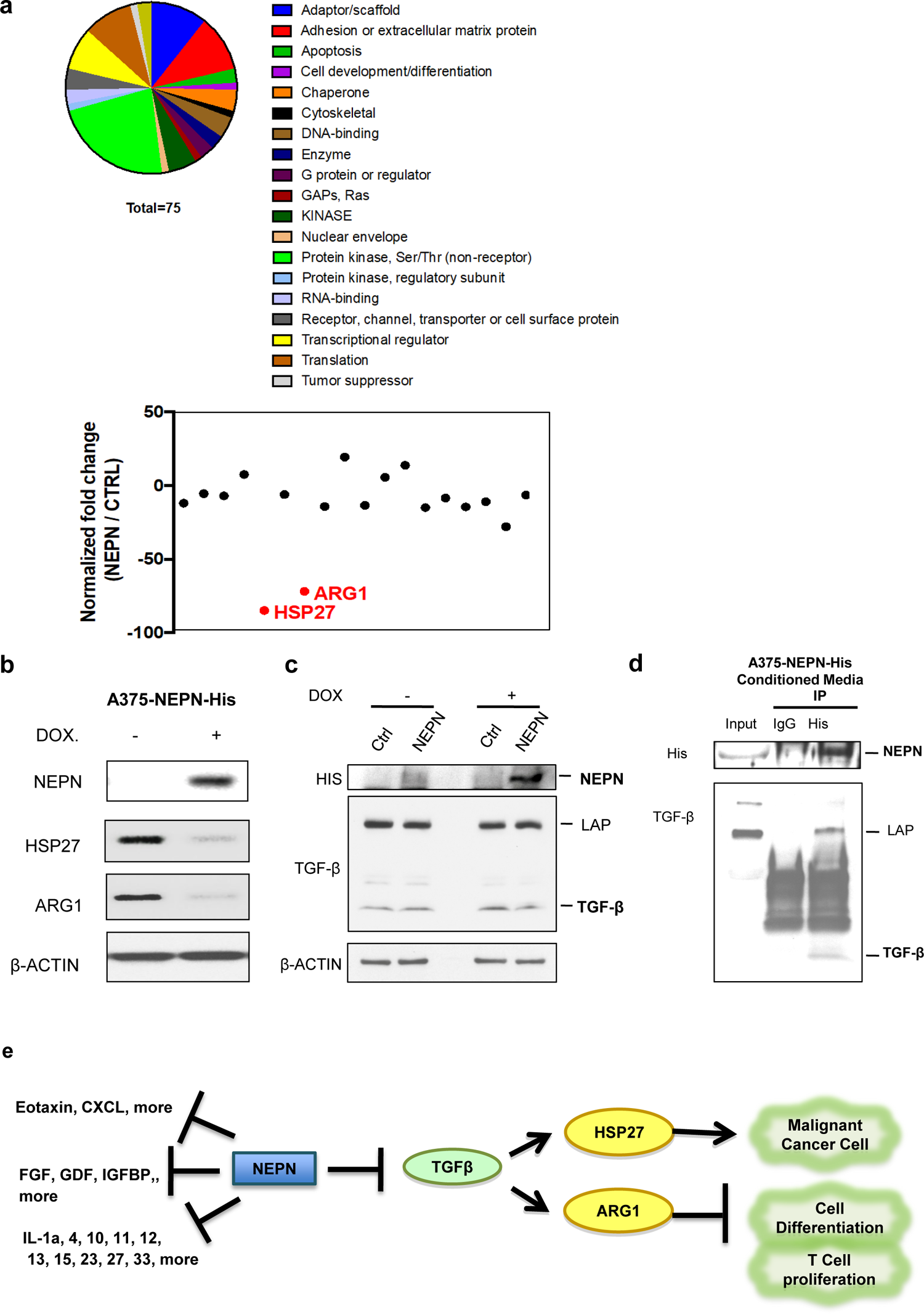
NEPN targets TGFβ signal pathway to suppress malignant progression. **a**, PTMScan-Mass Spectrometry direct proteomic profiling of downstream networks regulated by NEPN in A375 melanoma cells. The pie chart (top) illustrates NEPN-mediated signaling pathways based on quantification of 409 proteins and documentation of 1,006 unique phosphorylation and non-phosphorylation sites (n = 2, independent experiments). The dot plot (bottom) shows fold changes in protein expression and modification sites influenced by NEPN. **b**, HSP27 and ARG1 protein levels in A375 cells were confirmed by immunoblotting. **c**, NEPN expression reduces ARG1 and HSP27 but does not alter TGFβ1 protein levels. Western blot of A375 with and without NEPN expression with β-actin as loading control. **d,** Interaction of NEPN, TGFβ1 in conditioned medium was detected by co-immunoprecipitation. **e,** Hypothesis of NEPN mediated mechanism in cancer progression, including malignant progression, cell differentiation, T cell survival, and anti-tumor ability.

We confirmed that NEPN overexpression significantly reduces HSP27 and ARG1 protein levels in both melanoma and leukemia cells (**Fig. 3b and S.1f**) without significant effect on TGFb1 protein level (**Fig. 3c**). HSP27 is known to enhance the expression of metastatic genes that correlate with poor prognosis and to protect tumor cells from apoptosis and senescence.^23^ It also plays a key role in TGFβ1-induced epithelial-to-mesenchymal transition (EMT) *via* activation of MET kinase,^23^ which is a crucial event for cancer cells to acquire an invasive and metastatic phenotype.^28^ Additionally, HSP27 influences apoptosis in melanoma by favoring the ubiquitination of lκBα/p27^kip1^.^22,29^ ARG1, another downstream target of NEPN, has been reported to suppress T cell activation by two major pathways:^26^ (1) production of nitric oxide (NO; produced by ARG1 metabolism), which blocks the IL-2R signaling pathway and consequently prevents T cell proliferation; and (2) depletion of L-arginine in the local microenvironment, which results in impaired expression of the CD3ζ chain and inhibition of CD8+ T lymphocyte proliferation. And also, the TGFβ signal has been shown to increases ARG1 expression.^26^ Since NEPN belongs SLRP family which are key molecular effectors of cell surface and pericellular microenvironment (**Fig. 2c**),^9–11^ we hypothesized that NEPN inhibits HSP27 and ARG1 expression by blocking TGFβ signaling, either through sequestration of active TGFβ1 or interference with TGFβ receptor binding). Interestingly, both HSP27 and ARG1 are known effectors within the TGFβ cascade.^25,26^ By performing co-immunoprecipitation, we found that NEPN interacts with TGFβ complex included latent transforming growth factor β-binding protein (LTBP), TGFβ propeptide (LAP), and active TGFβ1, in melanoma and leukemia cells (**Fig. 3d and S1g**). These findings suggest that TGFβ and its associated proteins are key ligands within the NEPN-binding network, supporting a model in which NEM modulates tumor progression and immune evasion through direct inhibition of TGFβ1 signaling.

Aggressive cancer types, such as melanoma, are often highly resistant to standard treatment, including chemotherapy, targeted antibody inhibitors, and immunotherapies. As a result, patients frequently require combination therapies to overcome resistance and achieve more durable responses. TGFβ has emerged as a promising therapeutic target due to its well-established role as a tumor promoter, being abundantly expressed across multiple cancer types. Elevated levels of TGFβ correlate with enhanced tumor cell metastasis, while simultaneously suppressing host immune surveillance by modulating various immune cell lineages.^30–32^ In our model, the proliferation of various melanoma and leukemia cell lines was shown is strongly dependent on TGFβ1 signal pathway using TGFβ1 signal inhibitors, SB431542 and SB505734 (**Fig. S2a**). Additionally, inhibition of ARG1 using the inhibitors BEC and nor-NOHA also significantly reduced the proliferation rate of multiple human melanoma and leukemia cell lines (**Fig. S2b**). In further support, B16 melanoma cell line with a genetic knockout of ARG1 (ARG^KO^) displayed differentiated phenotypes with higher melanin contents and larger dark cell morphology (**Fig. S2c-d**). ARG1 inhibitors BEC and nor-NOHA also significantly inhibited the melanoma proliferation rate (**Fig. S2e**). Taken together, these findings highlight NEPN’s role as a novel anti-tumor factor that exerts its effects by modulating the TGFβ-ARG1 signal axis in tumor microenvironment (**Fig. 3e**).

### Broad Anti-Tumor Activity of NEPN Through TGFβ1 and Cytokine Regulation

In our *in vivo* B16 melanoma mouse model, NEPN overexpression led to a marked reduction in tumor growth signaling and significantly decreased tumor size, demonstrating the strong tumor inhibitory effect of NEPN (**Fig. 4a-b**). Furthermore, we observed a significant reduction in lung metastasis, indicating that NEPN also suppresses metastatic spread (**Fig. 4c**). To evaluate the therapeutic potential of purified NEPN protein in vivo, we expressed NEPN in 293 cells and purified it from the culture media using a His-tag (**Fig. S3f**). The resulting protein exhibited high stability, remaining intact after incubation at 37°C for 72 hours without detectable degradation (**Fig. 4d**). We conducted subcutaneous injections of B16-TGL melanoma cells, followed by localized systemic administration of purified NEPN protein once daily for 10 days (**Fig. 4e-g**). Remarkably, this short course of treatment cured 80% of mice (n = 10), which remained tumor-free and healthy through day 84, the end of the experiment, with no detectable toxic effects. In contrast, all untreated mice succumbed to tumor burden, with an average survival of only 35 days post-injection. These findings provide compelling evidence that NEPN functions as a potent and well-tolerated anti-tumor agent *in vivo* (**Fig. 4e-g**).

**Figure 4.**
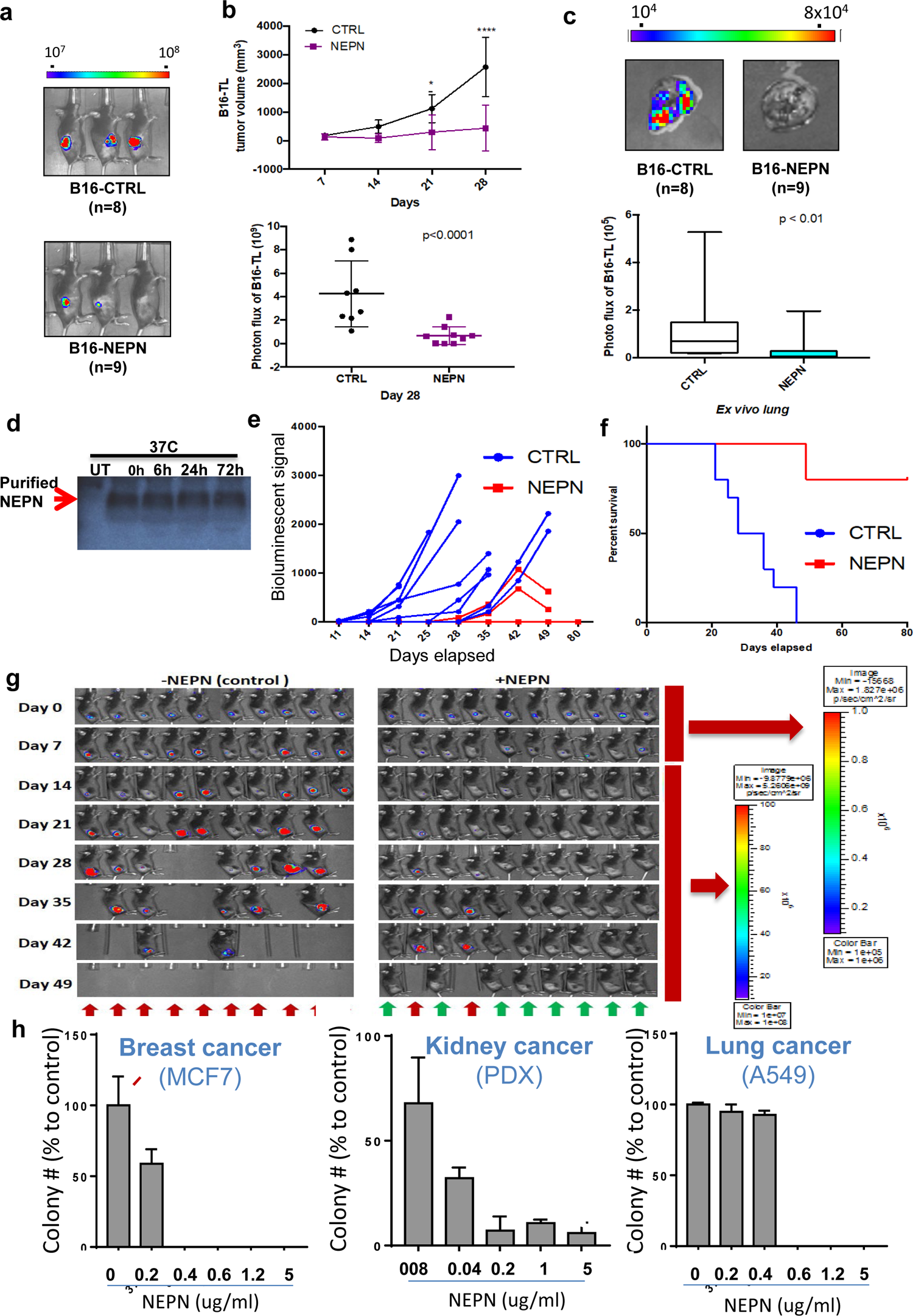
NEPN inhibits melanoma malignant progression. **a**, In vivo bioluminescent imaging of B16 melanoma cells (lentivirus-transduced with luciferase) with or without NEPN expression at day 28. Representative images are shown (n = 8-9, independent experiments). **b,** Tumor burden in vivo. Top: tumor volume was calculated weekly based on caliper measurements of tumor diameter (n = 8-9). Bottom: photon flux quantification of luciferase signal at day 28, confirming reduced tumor growth in NEPN-expressing tumors (n = 8-9). **c,** Ex vivo analysis of metastatic spread to the liver. Top: representative luciferase imaging of livers collected from tumor-bearing mice at day 28. Luminescence signals indicate metastatic B16 melanoma lesions. Bottom: quantification of liver photon flux shows significantly fewer metastases in NEPN-expressing tumors (CTRL n = 8, NEPN n = 9). **d,** Stability of purified NEPN protein. His-tag-purified NEPN expressed in 293 cells remained intact after incubation at 37 °C for 72 hours, with no detectable degradation. **e,** Mice bearing B16 melanoma tumors were treated with localized systemic administration of PBS or purified NEPN protein (5 µg) daily until day 11. Tumor size was measured weekly (bioluminescent signal). **f,** Kaplan-Meier survival analysis of control versus NEPN-treated mice (n = 10 biologically independent animals). **g,** Representative serial bioluminescence images of control (PBS) and NEPN-treated groups are shown from day 0 to day 49. Red arrows indicate mice that died, while green arrows indicate surviving mice with no detectable tumor signal through the end of the experiment (day 84). Note that day 0 and day 7 were imaged with 100-fold more sensitive luminescence settings than days 14-49. **h,** Broad-spectrum anti-tumor effects of NEPN across solid tumor models. Colony formation assays of MCF7 breast carcinoma, kidney cancer PDX (from the Department of Surgery, MSKCC, Rajasekhar Vinagolu), and A549 lung carcinoma cells treated with increasing concentrations of purified NEPN protein (0-5 µg/mL). NEPN reduced colony numbers in MCF7 and kidney PDX cells in a dose-dependent manner, with marked suppression at ≥0.2-0.4 µg/mL, and showed a sharp threshold response in A549 cells, with dramatic inhibition observed at ≥0.6 µg/mL. Data are representative of three independent experiments performed in triplicate. Statistical analysis: a-c, *P* values calculated using Mann-Whitney test. e, Gehan-Breslow-Wilcoxon test of survival of mice.

To determine whether NEPN’s anti-tumor activity extends beyond leukemia and melanoma, we evaluated its effects on colony formation in representative models of breast, kidney, and lung cancer (**Fig. 4h**). In MCF7 breast cancer cells, NEPN treatment resulted in a significant reduction in colony numbers in a dose-dependent manner, with nearly complete suppression at doses as low as 0.4 µg/ml. In a patient-derived xenograft (PDX) model of kidney cancer, NEPN also inhibited colony formation, with marked suppression observed at doses as low as 0.2 µg/ml. In contrast, In A549 lung cancer cells, NEPN showed a sharp threshold response: minimal change at 0-0.4 µg/ml, followed by dramatic, near-complete inhibition at ≥0.6 µg/mL. These findings demonstrate that NEPN possesses broad-spectrum anti-tumor activity, inhibiting the growth of breast, kidney, lung cancer cells in a dose-dependent fashion. Collectively, the data suggest that NEPN targets diverse malignancies and highlight its therapeutic potential across both hematologic and solid tumors.

### Reconstitution of human NEPN and cross-species anti-tumor activity

To develop a translational version of NEPN, we reconstituted the human NEPN sequence by aligning conserved domains from mouse and monkey NEPN with predicted human homologs. Evolutionary analysis revealed that although the NEPN protein is highly conserved across species, a key divergence occurred during primate evolution: when New World monkeys evolved into Old World monkeys (including humans), mutations in the N-terminal region led to deletion of the native secretory signal sequence and the introduction of premature stop codons.^33^ These alterations disrupted the natural secretion of NEPN, yet the core sequence remained highly conserved across primates, monkeys, and rodents, suggesting preservation of essential biological function.

To restore secretion and functionality, we corrected the premature stop codons and incorporated a humanized scFv signal peptide into the construct, enabling efficient secretion of human NEPN (hNEPN) (**Fig. S3a**). A His-tag was also included for purification. The engineered construct was cloned into a mammalian expression vector and expressed successfully, with robust secretion confirmed by biochemical assays.

Functional testing revealed that reconstituted hNEPN retained potent anti-tumor activity across species (**Fig. S3b**). In colony formation assays, hNEPN inhibited clonogenic growth of both human A375 melanoma cells and mouse B16 melanoma cells, demonstrating conserved cross-species efficacy. Dose-response analysis showed that A375 melanoma cells were sensitive to hNEPN at concentrations as low as 25 µg/mL, while B16 cells also exhibited significant inhibition. Direct comparison with mouse NEPN (mNEPN) confirmed that both variants suppressed A375 melanoma colonies, although mNEPN showed slightly stronger potency at equivalent doses. This observation suggests that further optimization of the N-terminal variable regions of hNEPN could enhance its anti-tumor activity.

Collectively, these results demonstrate that despite evolutionary mutations disrupting its secretion in humans, NEPN retains strong sequence conservation and functional potential, and engineered hNEPN can be reconstituted to exert tumor-suppressive effects. This provides a strong foundation for the development of NEPN-based therapeutic strategies.

### NEPN enhances the immune response and anti-tumor activity of T cells

The interactions among tumor cells, the microenvironment, and the immune system play a crucial in tumor progression. We hypothesize that NEPN inhibits tumor growth and metastasis not only directly by interrupting the ligand-receptor interactions, but also indirectly by enhancing T cell function. Given that ARG1 is closely linked to T cell activation and immune regulation, it is likely that NEPN’s anti-tumor effects are partially dependent on modulating the immune system to support its function.

We designed an *in vivo* experiment to evaluate the therapeutic effects of NEPN in immunocompetent (B6) (**Fig. S4a-c)** and immune-deficient (Rag2gammaC) (**Fig. S4d-f)** mouse models with a suboptimal dose (20% of the established effective dose). Interestingly, NEPN significantly suppresses melanoma progression in B6 mice but has a lower effect on Rag2gammaC mice. These results suggested that NEPN’s anti-tumor function partially depends on an intact host immune system. Furthermore, a complete blood count assay revealed increased levels of white blood cells (WBC), neutrophils (NEU), lymphocytes (LYM), and eosinophils (EOS) in treated host after 7 days. This suggests that NEPN might enhance the immune response within the tumor microenvironment, potentially involving T cells, dendritic cells and macrophage (data not shown).

Using an *in vivo* T cell transplantation model, we demonstrated that NEPN enhances human T cell proliferation in a time-dependent manner (**Fig. 5a**). This proliferative effect was consistently observed across experiments involving three different donors (**Fig. 5b**). Based on our observation, we propose two key mechanisms underlying NEPN’s enhancement of T cell function: (1) NEPN-mediated inhibition of ARG1 likely increases local arginine concentrations, a crucial regulator known to promote T cell survival and anti-tumor activity;^34^ (2) TGFβ is a pivotal regulator of thymic T cell development, peripheral T cell homeostasis, tolerance to self-antigens, and T cell differentiation.^35–38^ By blocking of TGFβ1 activity within the tumor microenvironment, NEPN may preferentially support the differentiation of effector T cells while limiting regulatory T cell development, therefore enhancing anti-tumor immune responses.^35–38^

**Figure 5.**
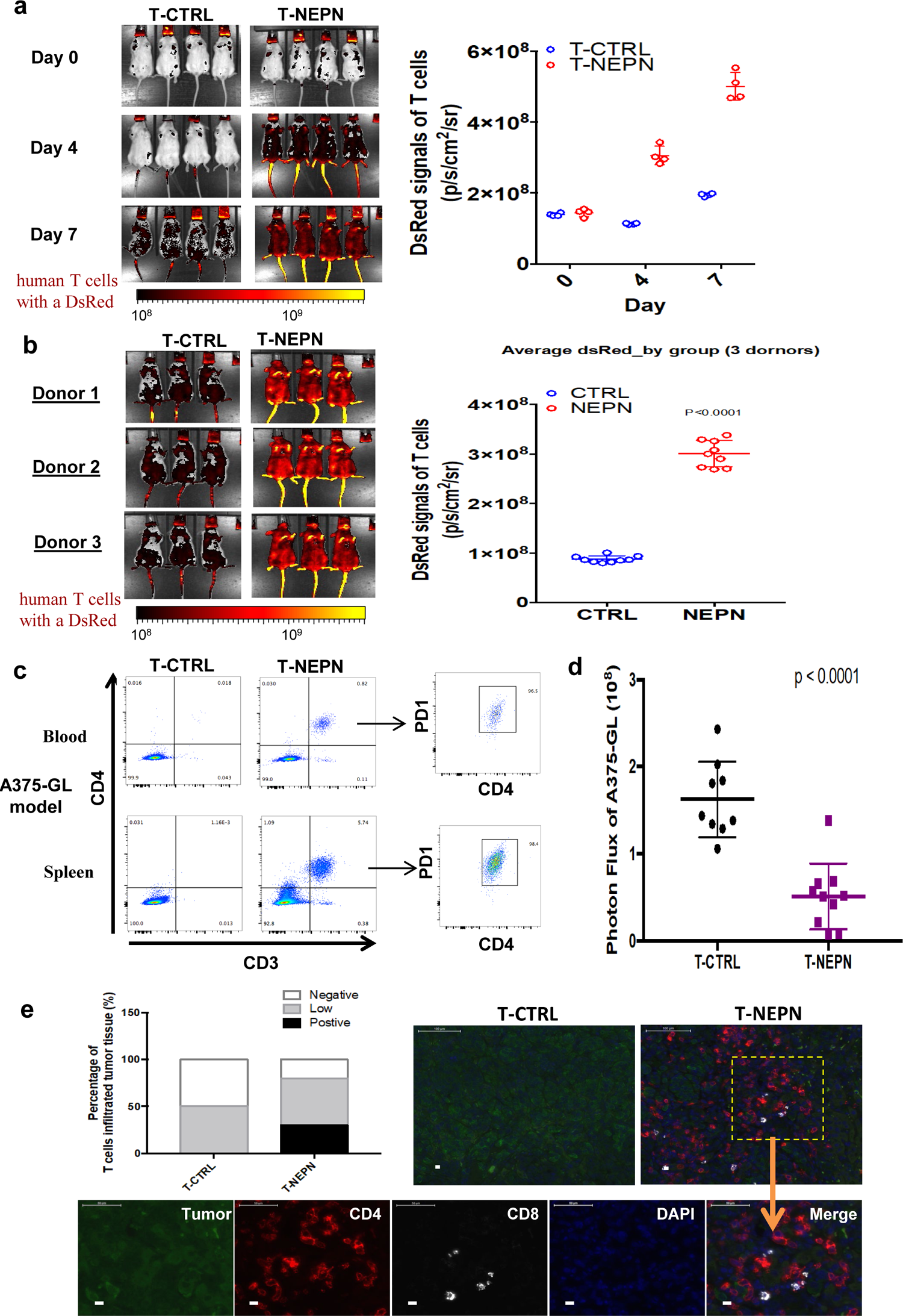
NEPN enhances the antitumor activity of T cells. **a** Proliferation signal of T-cells by measuring DsRed signal (n = 4 biologically independent). T-Test. **b,** Effect of NEPN on T cell proliferation from 3 different donors (n = 9, 3 independent mice on 3 different donors). **c,** Flow cytometry analysis of peripheral T cell population from blood or spleen on day 18 with the antibodies specific to CD3, CD4, and PD1. **d,** Bioluminescent signals of melanoma A375-TGL on day 7 *in vivo* xenograft model (n = 10, biological independent). **e,** Tumor-infiltrating T cells in the melanoma model. Immunohistochemistry of CD4⁺ and CD8⁺ T cells in melanoma tumors (n = 10). Tumor infiltration was classified as negative (<5% T cells/field), low (5-20%), or positive (>20%). Bar graph shows the distribution of infiltration levels in control versus NEPN-treated groups. Representative images show tumor tissue (green), CD4 (red), CD8 (white), DAPI (blue), and merged views. Scale bars, 100 μm. a-c, *P* values calculated using Mann-Whitney test.

For clinical applications, we developed a novel engraftment model combining T cell-based NEPN therapy^39,40^ to efficiently deliver NEPN to the tumor microenvironment alongside T cell therapy to better control tumor progression. Enhanced T cell activation and proliferation were observed in the NEPN-treated group (**Fig. S5a-b**), with a particularly notable increase in CD4^+^ T cells (**Fig. 5c** and **Fig.S5c**). Tumor burden was significantly delayed in both melanoma and leukemia models (**Fig. 5d** and **Fig.S5d**). Furthermore, analysis of tumor-infiltrating lymphocytes revealed a significant increase in CD4^+^ and CD8^+^ cells at the liver metastasis site in the melanoma model (**Fig. 5e)**. Given that elevated TGFβ levels are associated with reduced tumor infiltrated by cytotoxic T cells ^41^, these findings support that NEPN captures TGFβ and modulate the TME. Collectively, these data suggest NEPN not only inhibits tumorigenesis and metastasis but also enhances T cell activity and host immune surveillance capacity.

### Immunological delivery of NEPN by CART cells enhances tumor suppression and reduces relapses

To establish a pre-clinical model for NEPN delivery into the tumor microenvironment, we employed an immunological approach using engineered chimeric antigen receptor (CAR) T cells. CART therapy is a clinically established immunotherapy in which patient-derived T cells are modified to express synthetic receptors containing an antibody-derived extracellular domain coupled to intracellular T cell signaling modules such as CD28 and CD3ζ. This hybrid design enables CART cells to specifically recognize tumor-associated antigens and trigger T cell activation. By engineering CART cells to express NEPN, we sought to combine direct tumor targeting with localized release of NEPN to potentiate immune-mediated tumor killing.

For proof-of-principle, we utilized the clinically verified 1928Z CAR construct, which targets CD19⁺ leukemia cells and has been widely validated in translational studies (**Fig. 6a**). NALM6 leukemia cells were injected on day-4, followed by intravenous infusion of either control CART cells or NEPN-expressing CART cells on day 0. Bioluminescence imaging on day 4 demonstrated that untreated mice displayed rapid leukemia progression, whereas NEPN-CART-treated mice exhibited significantly lower tumor burden compared to both CART alone and control groups (**Fig. 6b**). Tumor growth kinetics revealed that, at a suboptimal CART dose of 0.1 million cells, NEPN-CART therapy enhanced tumor clearance even within the first four days post-treatment (**Fig. 6b**) and reduced relapse events up to day 35 (**Fig. 6c**), achieving superior early suppression relative to CART cells alone. At a higher CART dose of 0.2 million cells, NEPN-CART treatment sustained tumor control and markedly reduced relapse events up to day 35, whereas tumors in the CART-only group relapsed more frequently (**Fig. 6c**).

**Figure 6.**
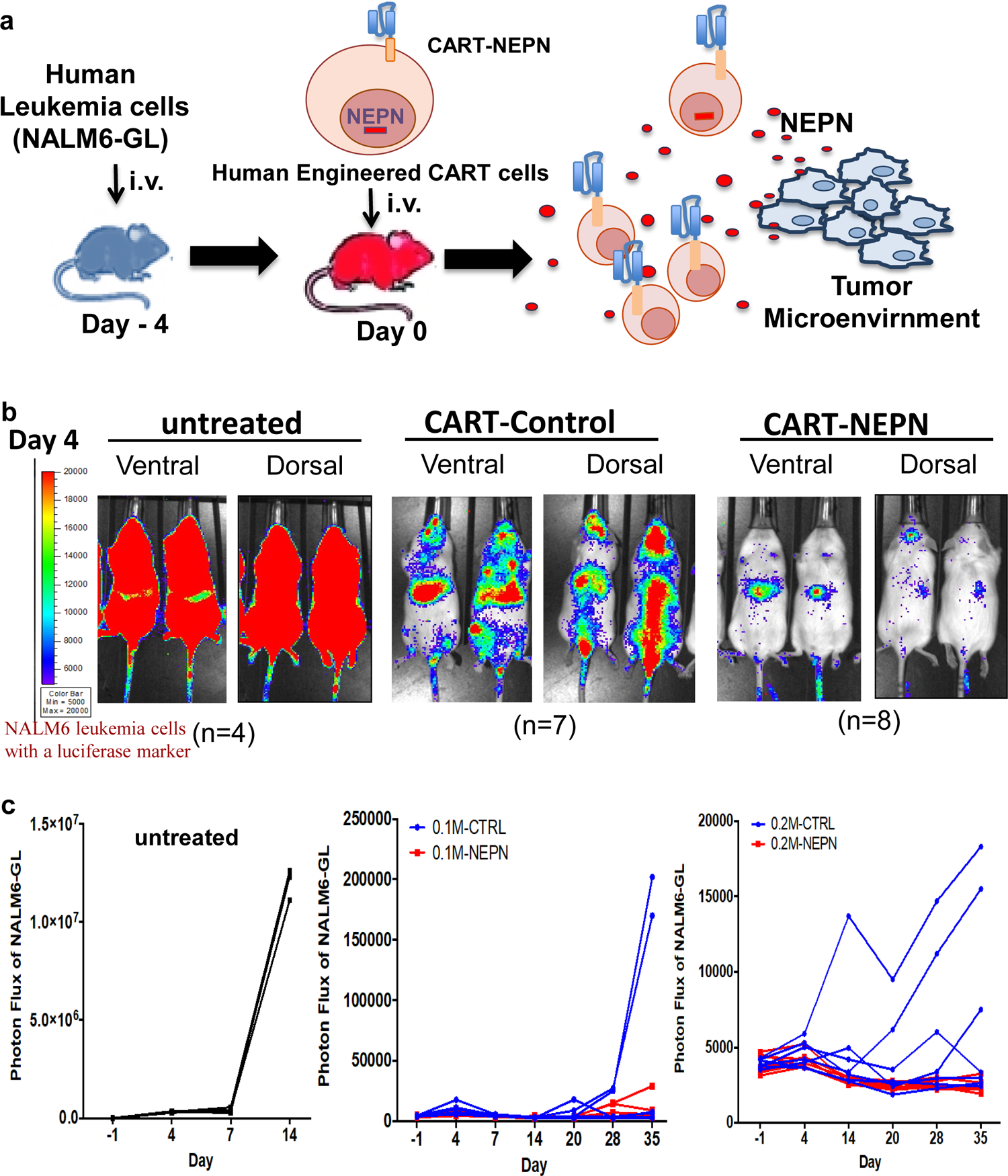
Immunological delivery of NEPN by CART cells enhances tumor suppression. **a**, Schematic of experimental workflow: NALM6 leukemia cells (with a luciferase marker) were injected on day-4, followed by infusion of control CART or NEPN-expressing CART cells (1928Z, anti-CD19) on day 0. Engineered CART cells target CD19⁺ tumor cells and release NEPN into the tumor microenvironment. **b,** Representative bioluminescence images showing untreated mice, 0.1 million of CART-treated mice and NEPN-CART-treated mice. NEPN-CART treatment led to stronger tumor suppression than CART alone. **c,** Tumor growth kinetics measured by photon flux. Untreated tumors (n=4) progressed rapidly, while NEPN-CART treatment showed enhanced tumor killing at both 0.1 million (n=7) and 0.2 million (n=8) CART doses (suboptimal level). NEPN-CART-treated mice exhibited reduced relapse and sustained tumor control up to day 35 compared to CART alone.

Collectively, these results indicate that CART cells engineered to deliver NEPN act synergistically within the tumor microenvironment, producing stronger tumor suppression and depressing relapse compared to conventional CART therapy. This strategy highlights NEPN as a potent adjunct for adoptive cell immunotherapy.

### Define the ligand-receptor network of NEPN-mediated mechanism on malignant tumor growth and metastasis

We hypothesize that NEPN controls tumor growth and death by modifying the extracellular environment and regulating key cellular signaling pathways. Post-translational modification profiling analysis (PTMScan) identified HSP27 (inhibits apoptotic pathway)^22,23,29^ and ARG1 (inhibits T cell activation)^24–26^ are the most significantly influenced downstream targets of NEPN (**Fig. 3a**). Notably, both proteins share a common upstream TGFβ regulatory pathway.^26,27^ We propose that NEPN inhibits HSP27 and ARG1 by blocking ligands-receptor signals such as TGFβ either through sequestration of ligands or blocking of receptors), thereby suppressing tumor formation and metastasis (**Fig. 3e**). To further define the molecular partners of NEPN, we performed immunoprecipitation coupled to mass spectrometry (IP-MS) in tumor lysates. Comparative analysis with two independent controls (CTRL1 and CTRL2) identified 100 proteins that were specifically enriched in both NEPN replicates (NEPN1 and NEPN2) but absent in controls (**Fig. S6a-c**). These candidates included ribosomal proteins (Rpl family), RNA-binding proteins (Snrnp, Hnrnp families), cytoskeletal components (Krt, Tsr), metabolic regulators (Aldh5a1, Gpx4), and signaling molecules (Rasa3, Serping1), indicating broad interaction of NEPN with structural and signaling pathways. Importantly, LRBP was identified among the interactors, representing a strong candidate in the ligand-receptor binding network that connects NEPN to upstream cytokine signaling. These findings support our hypothesis that NEPN exerts its tumor-suppressive activity by modulating ligand-receptor interactions and thereby regulating downstream effectors such as HSP27 and ARG1. (**Fig. S6**).

Additionally, we also identified 44 cytokines that interacted with NEPN using an immunoprecipitation-cytokine array (**Fig. S7a-b**). Pathway enrichment analysis showed that NEPN is involved in cytokine-cytokine receptor interaction, chemokine signaling, T cell receptor signaling, immune response, extracellular matrix organization, and other important pathways (**Fig. S7c**). For example, the Th2 cytokine IL-4 and IL-13 are related to cancer aggressiveness^42^ and IL-4 blockade has been shown to enhance response to immunotherapy.^43^ Moreover, NEPN promotes the chemotaxis of T cells, B cells, and monocytes, and natural killer cells. In addition, both IL-10 and TGF-β are known to cooperate in the induction and maintenance of regulatory T cells (Tregs), which play a central role in establishing immune suppression within the tumor microenvironment. Our findings raise the possibility that NEPN may modulate this axis by altering cytokine availability or signaling, thereby impacting Treg activity and broader immune balance. This regulatory T cell-mediated mechanism will be further investigated in future studies.^44^ Through the immune modulation in the tumor microenvironment, it might change the phenotype of tumor cells and immune cell phenotypes. To assess the clinical value of NEPN, establishing a functional humanized immune system is critical to investigate NEPN’s role at the interface cancer cells and the immune system within the tumor microenvironment. Our works reveals NEPN as a novel anti-tumor factor originating from the early embryonic stage with promising therapeutic potential cancer immunotherapy. We have demonstrated that NEPN modulates the tumor microenvironment to both suppress tumor growth and enhance immune responses. We believe the most effective therapeutic strategy combines tumor control with immune activation, and NEPM embodies this dual function.

## METHODS

### Cell culture and reagent

Cells were maintained at 37°C in DMEM (supplemented with 10% heat-inactivated fetal bovine serum (FBS) and 2 mM L-glutamine, 100 U/ml penicillin and 100 μg/ml streptomycin) or RPMI-1640 medium (modified to contain 2 mM L-glutamine, 10 mM HEPES, 1 mM sodium pyruvate, 4500 mg/L glucose, and 1500 mg/L sodium bicarbonate), in a humidified atmosphere with 5% CO_2_. Cell viability was measured by using trypan blue exclusion. Mouse melanoma B16, human melanoma A375 and derived cell lines were cultured in a complete DMEM medium. Human uveal melanoma L3OMM1.3, PDX cell line Mel1, human B cell precursor leukemia NALM6 cell line, and derived cell lines were cultured in RPMI-1640 complete medium.

### Co-culture assay

Embryonic skin cells 1×10^4^ E14.5-NEPN were plated in 6 well plates with or without doxycycline and allowed to adhere overnight. The next day, 2×10^4^ B16 melanoma-Cyan cells were seeded and co-cultured for 72 hr. The conditional medium was collected 72 hr after culturing primary embryonic tissues and used to treat melanoma cells for colony formation assay.

### Colony formation, migration, and invasion assay

50-250 cells were seed to each well of a 48-well or 24-well culture plate and incubated for 7 days at 37 ℃. The colonies are fixed by 100% methanol at room temperature for 20 min, following stained using crystal violet staining solution for 5 min.^45^ The images were analyzed using Image J software. Migration and invasion assays were based on the Boyden chamber transwell model. Migration assays were performed using 8-μm pore polyester transwell inserts (Corning, MA) equilibrated in PBS buffer. Cells were pre-cultured in serum-starved medium (0.2%FBS, overnight), re-suspended (5×10^4^ cells) in 100 μl of serum-free medium, and added to the upper chamber. A culture medium containing 10% FBS was added to the lower chamber. The cellular migration was allowed to proceed for 6 hours at 37°C in a 5% CO_2_ incubator. For invasion assays, the inserts were pre-coated with matrigel for 3 hours and the cellular invasion was allowed to proceed for 24 hours. The inserts were then removed, washed with PBS, fixed 10 minutes in 100% methanol, and stained for 30 minutes by using 2% crystal violet. Migrated and Invasive cells were counted in five random fields.

### PTMScan direct proteomic profiling analysis

Human A375-NEPN were cultured for 7 days with or without doxycycline and protein lysates were prepared according to the sample preparation protocol for PTMScan^®^ Analysis (Cell Singling). Samples were processed and the protein candidates were identified by Cell Signaling Technology, Inc.

### RNAseq and real-time quantitative PCR

RNA was extracted from embryonic tissues using RNeasy® Micro Kit (Qiagen) or cells using RNeasy® Mini Kit (Qiagen), following the manufacturer’s instructions. For RNAseq analysis, the RNA was sequencing and analyzed by the IGO core facility (MSKCC). For real-time quantitative PCR, the procedure is according to the previous publication^46^. Briefly, total RNA was reverse transcribed to cDNA using the M-MuLV reverse transcriptase system (New England Biolabs). The expression levels of interesting genes (Nepn, Habp2, and β-actin) were quantified by real-time qPCR reaction using the Power SYBR Green PCR Mastermix (Applied Biosystems). The conditions were programmed as follows: initial denaturation at 95 °C for 10 min followed by 40 cycles of 30 s at 95 °C, 1 min at 55 °C and 1 min at 72 °C; then 1 min at 95 °C, 30 s at 55 °C and 30 s at 95 °C. All of the samples were duplicated, and the PCR reaction was carried out using a Mx3005P reader (Stratagene). The amount of PCR product was normalized as a percentage of the expression level of β-actin. The sequences of the primers used are provided in the Supplementary Table.

### Melanin Detection Assay

Cells were incubated in 1N NaOH to solubilized melanin. Spectrophotometric analysis of the melanin content was then conducted at 405 nm; the total protein concentration of the cells was determined using a BCA protein assay kit. The melanin content was determined based on the absorbance/ug of the protein^47^.

### Immunoprecipitation and Immunoblotting

Treated and untreated cells were collected and performed immunoprecipitation using PureProtemone^TM^ Nickel Magnetic Beads according to the user guide.

Briefly, a 2ml condition medium was incubated with 200 μl of magnetic bead suspension in Pierce IP Lysis Buffer overnight, followed by a wash and elution step. Protein concentrations were determined using a BCA protein assay kit (Pierce) and immunoblotting^46^ is performed. Antibody information is provided in the Supplementary Table.

### Cytokine Array

Samples after immunoprecipitation continue the procedure using mouse XL Cytokine Array Kit (R&D Systems, Inc.), following the manufacturer’s instructions. Briefly, immunoprecipitated proteins were dissolved in PBS with 1% Triton X-100 and protease inhibitors. The array membranes were incubated with immunoprecipitated samples overnight at 4°C, followed incubated with detection antibody cocktail for 1 hr and 1X Streptavidin-HRP for 30 min at room temperature. The membranes were reacted with Chemi Reagent Mix and exposed to X-ray film.

### Lentivirus production

293T cells were transfected with pDOX-CTRL-tdTomatoLuc, pDOX-NEPN-dTomatoLuc, pLenti-RTTA using calcium phosphate cell transfection, following the virus collection and concentration as previously described^46^.

### Retrovirus Production

NEPN-DsREd or DsRed were cloning into the SFG γ-retroviral (RV) vector^48^. VSV-G pseudotyped retroviral supernatants derived from transduced gpg29 fibroblasts (H29) were used to construct stable retroviral-producing cell lines as previously described^49^. T cells were transduced with virus supernatants using centrifugation on retronectin (Takara)-coated plates.

### CRISPR knockout

pLentiCRISPR v2 vector with mouse Arg1 CRISPR ((LV441107)) guide RNA and 5 or human ARG1 CRISPR (LV078906) RNA 2 and 3 were purchased from GenScript. Melanoma cells were infected and the targeted colonies were drug-selected using puromycin and protein expression levels were analyzed using immunoblotting.

### Antibodies and cell staining

For the flow cytometry analysis, cells were stained with the indicated antibodies and live-dead cell staining dye. Information of the antibodies and staining dye is provided in Supplementary Table.

### Protein purification

293T cells were infected pDOX-NEPN-His-tdTomato lentivirus and sorted by tdTomato signaling. The culture medium was collected 72 hr after doxycycline treatment and secreted NEPN-His protein was purified using Capturem^TM^ his-tagged purification maxiprep columns (Takara) and concentrated using Amicon Ultra-15 centrifugal filter units, following the manufacturer’s instructions. Purified protein was detected using coomassie blue staining and glycoprotein staining Kit (Pierce).

### Immunohistochemistry

Tissue was collected from mice and fixed in formalin solution 10%, neutral buffer. Immunohistochemistry staining was performed by the Molecular Cytology facility (MSKCC).

### Colony Formation Assays in Breast, Kidney, and Lung Cancer Models

MCF7 breast carcinoma and A549 lung carcinoma cell lines (ATCC) were maintained in DMEM supplemented with 10% FBS, 1% penicillin-streptomycin, and 2 mM L-glutamine at 37 °C with 5% CO₂, while kidney cancer PDX cells were obtained from the Department of Surgery, Memorial Sloan Kettering Cancer Center (Rajasekhar Vinagolu) and cultured in RPMI-1640 with 10% FBS. All cell lines were authenticated by STR profiling and confirmed mycoplasma-free. Purified NEPN protein was expressed in HEK293 cells using a His-tag construct and isolated from culture media by affinity chromatography. For colony formation assays, cells were seeded at low density (200-500 cells/well in 6-well plates) and treated with increasing concentrations of NEPN (0-5 µg/mL). Medium was replaced every 3 days with fresh NEPN-containing media, and colonies were allowed to grow for 10-14 days before fixation with methanol, crystal violet staining, and quantification by manual counting or ImageJ. Colony numbers were normalized to untreated controls, and data were analyzed from three independent experiments in triplicate using one-way ANOVA with Dunnett’s post-hoc test, with P < 0.05 considered significant.

### Animal studies

All experiments using animals were performed under our protocol approved by MSKCC Institutional Animal Care and Use Committee (IACUC). 8- to 12-week-old C57BL/6JB6 and NOD/SCID/IL-2Rγ -\ null (NSG) female mice (Jackson Laboratory) and Rag2/Il2rg double knockout mice (Taconic) were used for our melanoma models. NOD/SCID/IL-2Rγ -\ null (NSG) male mice were used for our leukemia model.

For melanoma models, mice were intracardiac or subcutaneous injected into recipient mice. Mice were injected with 0.2 M melanoma cells and subcutaneously injected with purified NEPN protein or PBS into the tumor location daily until day 14. For the T-cell delivery model, mice were injected with engineered T cells 4 days after tumor implantation.

For leukemia models, mice were inoculated with 0.5M NALM6-GL (GFP-luciferase) cells via tail vein, followed by T cell or CART cell injected four days later.

Bioluminescence and fluorescence imaging detected using the Xenogen IVIS Imagining System (Caliper Life Sciences) and analyzed using Living Image software (Xengen). Tumors volume was measured and calculated using this formula: (D X d^2^)/2, in which D and d refer to the long and short tumor diameter, respectively.

### Generation and Evaluation of NEPN-Engineered CART Cells in a NALM6 Xenograft Model

NALM6 leukemia cells expressing GFP and firefly luciferase (NALM6-GL) were cultured in RPMI-1640 with 10% FBS, 1% penicillin-streptomycin, and 2 mM L-glutamine at 37 °C, 5% CO₂, and confirmed mycoplasma-free. Primary human T cells were isolated from PBMCs, CD3⁺ enriched, and activated with anti-CD3/CD28 Dynabeads plus IL-2 (50 IU/mL) before lentiviral transduction with the clinically validated CD19-targeting 1928ζ CAR (scFv-CD19/CD28/CD3ζ). For NEPN-CART cells, NEPN was cloned downstream of a P2A sequence to enable bicistronic CAR and NEPN secretion; expression was verified by immunoblotting and ELISA, and function by T cell-tumor co-culture. Six- to eight-week-old NSG mice (The Jackson Laboratory) were injected i.v. with 2 × 10⁵ NALM6-GL cells (day-4), followed by 0.1 × 10⁶ or 0.2 × 10⁶ CART or NEPN-CART cells (day 0); controls included untreated mice. Tumor burden was monitored weekly by IVIS bioluminescence after D-luciferin injection (150 mg/kg), quantified as photon flux using Living Image software. Tumor growth was analyzed by Mann-Whitney U test and survival by Gehan-Breslow-Wilcoxon test in GraphPad Prism v9, with P < 0.05 considered significant.

## Ethics / Approvals

All animal experiments were conducted in accordance with protocols approved by the Institutional Animal Care and Use Committee (IACUC).

## Author Contributions / Acknowledgments / Funding

K.K. conceived, supervised, and wrote the project and manuscript. T.C. led the project, performed experiments, analyzed data, and contributed to manuscript writing. Y.H., X.H., N.K., R.V., and C.Z. performed experiments and assisted with data analysis. C.Z. conducted computational analysis. R.V., R.Z., S.M.F., H.L., S.L., and M.S. provided supervision and project consultation.

## Licensing / Reuse Statement

This preprint is made available under a CC-BY 4.0 International license, which permits unrestricted use, distribution, and reproduction in any medium, provided the original work is properly cited.

---

**Supplementary Figure 1.**
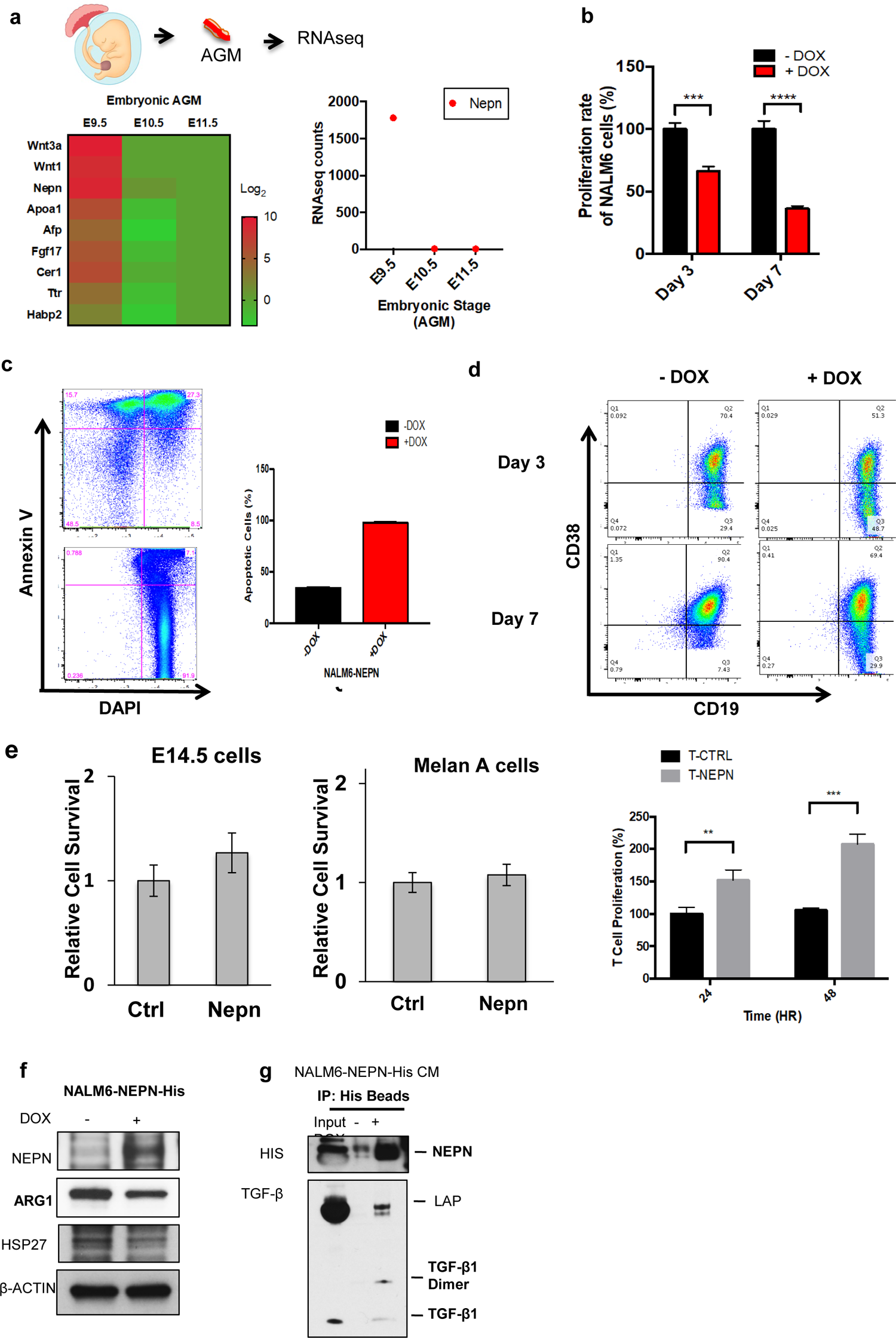
**a**, Heatmap (left) and counts (right) of gene expression pattern in AGM region at different embryonic stages using RNAseq analysis (n=2, independent experiments). **b,** Proliferation rate of NALM6-NEPN after doxycycline induction. **c,** Apoptosis of NALM6-NEPN cells after doxycycline induction. Measured by Annexin V/PI staining and flow cytometry (Annexin V-FITC Kit, BD Biosciences) in 48 h. Representative plots (left) and quantification from three independent replicates (right) show increased apoptosis with +DOX compared to - DOX. **d,** Differentiation of NALM6-NEPN cells following NEPN induction. Flow cytometry analysis of CD19 and CD38 expression in NALM6-NEPN cells at day 3 (top) and day 7 (bottom) with or without doxycycline induction. NEPN expression increased the proportion of CD19⁺CD38⁺ differentiated populations, indicating that NEPN promotes leukemia cell differentiation over time. **e,** The effect of NEPN on normal cells. Relative survival of E14.5 embryonic skin cells and Melan-A melanocytes was measured by Trypan Blue exclusion/automated viable cell counting, normalized to - DOX controls in 48 h; no cytotoxicity with NEPN induction was observed. Primary human T cells were quantified by direct viable cell counts at 24 and 48 h after NEPN induction, showing increased proliferation versus T-CTRL. **f**, HSP27, and ARG1 protein levels in NALM6 cells were confirmed by immunoblotting. **g**, The interaction of NEPN, TGFβ1, and TGFβ receptor in NALM6-NEPN derived conditioned medium (CM) was detected by co-immunoprecipitation. * P < 0.05, * * P < 0.01, * * * P < 0.001, * * * * P < 0.0001(t-test).

**Supplementary Figure 2.**
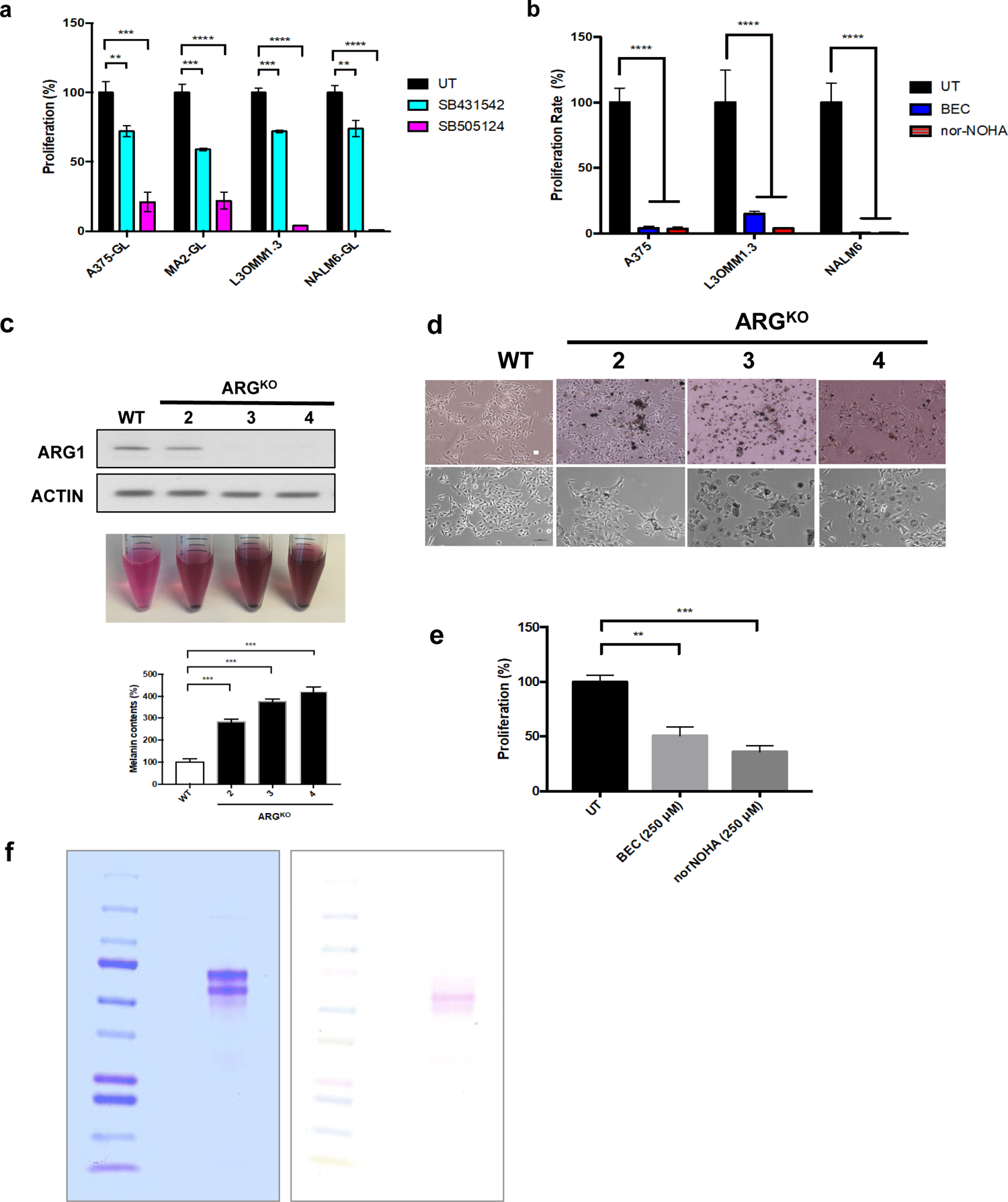
**a**, Proliferation rate of different human melanoma and leukemia cells treated with TGFβ inhibitors, SB431542 or SB505124. **b,** the Proliferation rate of different melanoma and leukemia cells treated with 250uM arginase inhibitors, BEC or nor-NOHA, by measuring luciferase signal. **c,** ARG1 protein level of B16 and B16-ARG^KO^ cell lines were detected using immunoblotting (top). Melanin level in these cell lines was detected (bottom). **d,** The phenotype of B16 and B16-ARG^KO^ cells. Scale bar, 100 μm. **e,** Proliferation rate of B16 cell treated with 250nM arginase inhibitors, BEC or norNOHA. * P < 0.05, * * P < 0.01, * * * P < 0.001, * * * * P < 0.0001(t-test). **f,** The purity of the purified mouse NEPN using SDS gel, following Coomassie blue staining (left) and glycoprotein staining (right).

**Supplementary Figure 3.**
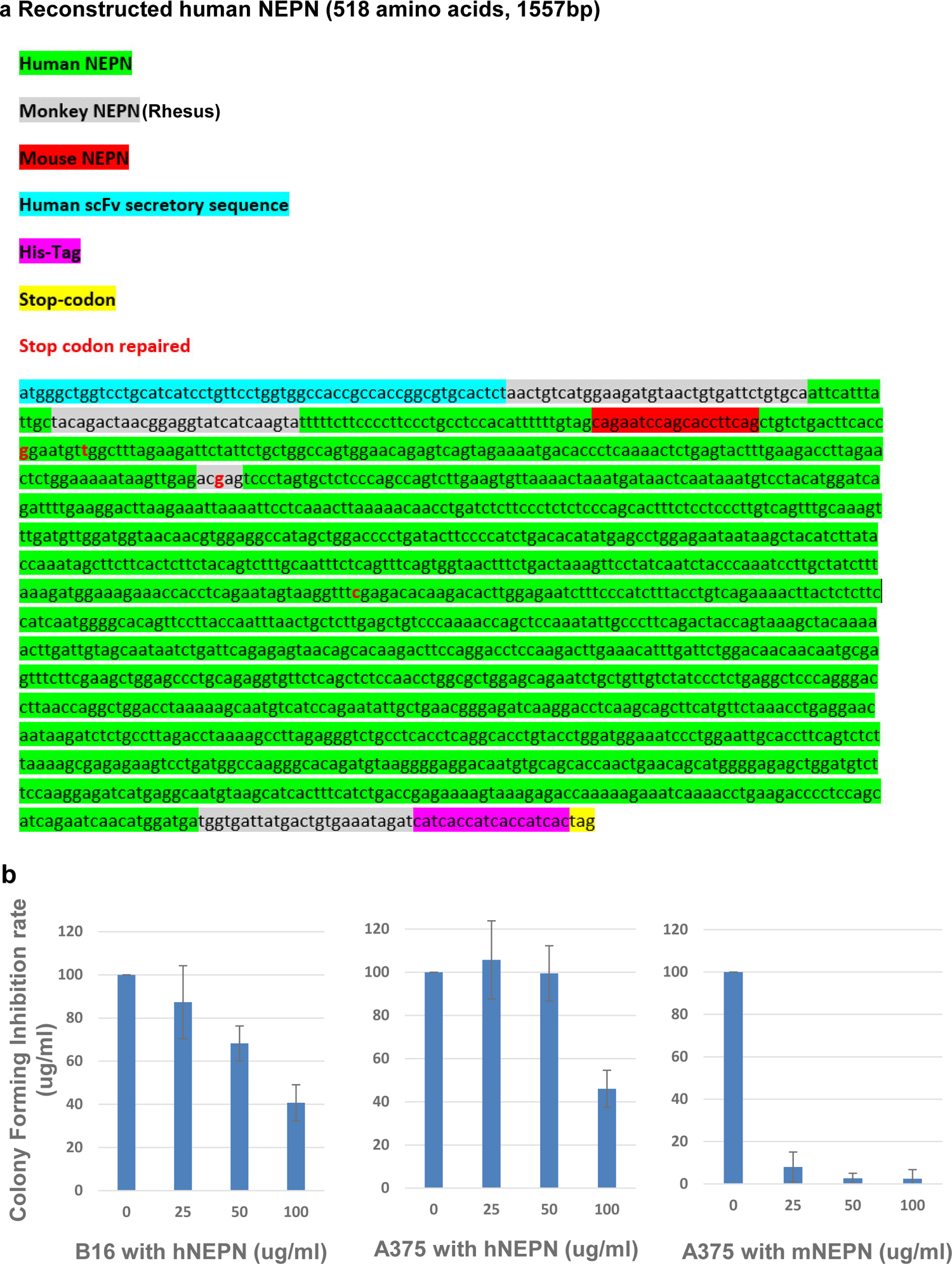
Reconstitution and functional testing of human NEPN. **a,** Schematic sequence map of reconstituted human NEPN showing conserved coding regions across species, repaired stop codon, addition of human scFv secretion signal, and His-tag. The monkey NEPN DNA sequence used primarily derived from rhesus macaque, with comparative analysis performed across 12 old-world and new-world monkey species. This revealed that the intact NEPN gene is conserved in lineages closer to humans. **b,** Colony formation assays with reconstituted human NEPN (hNEPN) in B16 (mouse) and A375 (human) melanoma cells show dose-dependent inhibition of colony formation. Mouse NEPN (mNEPN) served as control in A375 cells, demonstrating stronger potency compared to hNEPN. These results confirm cross-species activity of NEPN and highlight the need for optimization of the variable N-terminal regions of human NEPN to enhance therapeutic potency.

**Supplementary Figure 4.**
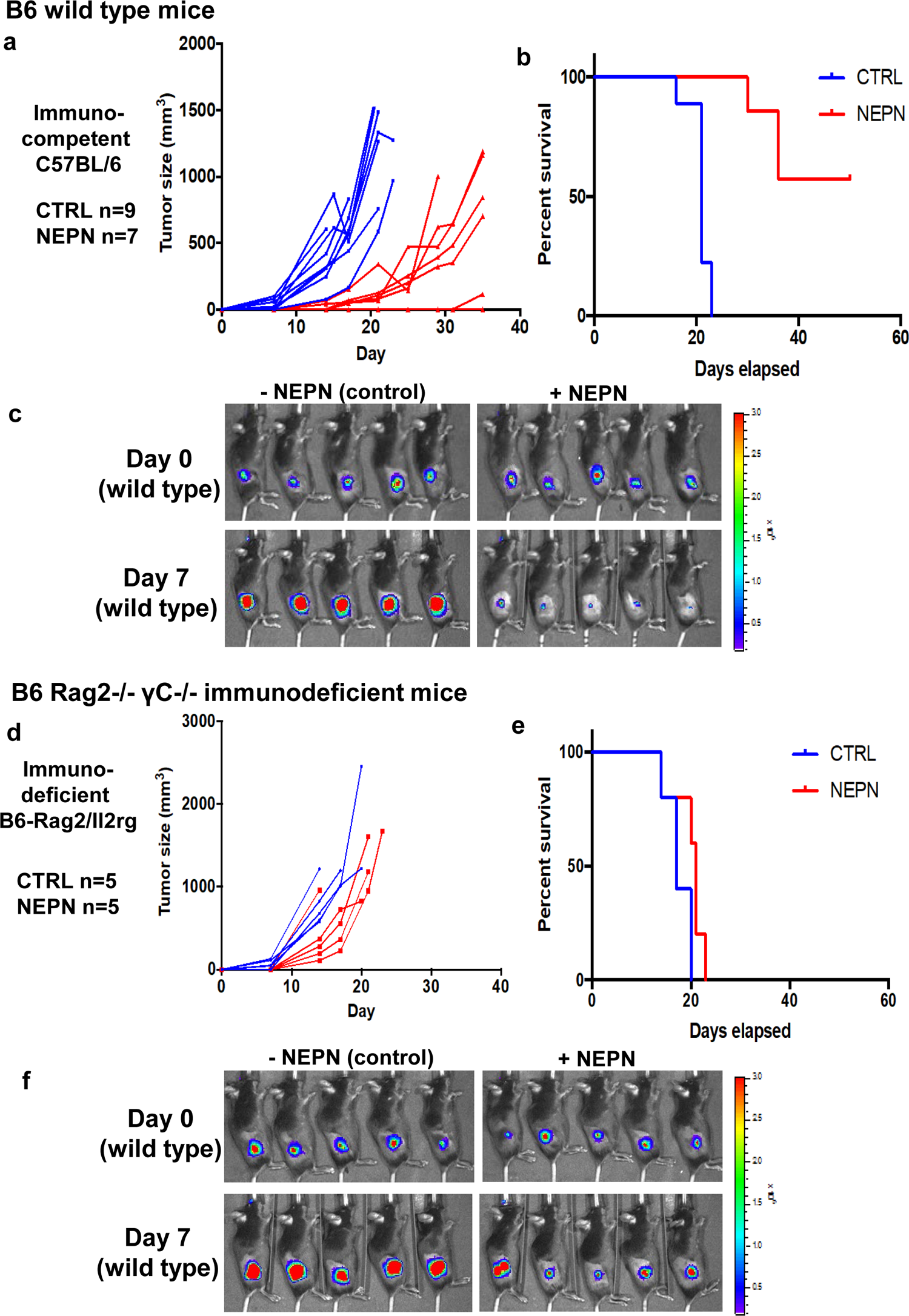
Suppression of melanoma growth by purified NEPN injection in immunocompetent and immunodeficient mice. 200,000 B16 melanoma cells were injected subcutaneously, and mice received daily injections of purified NEPN protein (1 µg, 20% of optimal dose) or PBS control from day 1 to day 11. Tumor growth was monitored by size and bioluminescent imaging, and survival was recorded. **a,** Tumor growth curves in C57BL/6 (B6, immunocompetent) mice treated with NEPN or PBS. **b,** Kaplan-Meier survival analysis of B6 mice. **c,** Representative bioluminescence images of B6 mice at day 7 showing reduced tumor burden in NEPN-treated animals. **d,** Tumor growth curves in B6-Rag2/Il2rg (Rag2γc, immune-deficient) mice treated with NEPN or PBS. **e,** Kaplan-Meier survival analysis of Rag2γc mice. **f,** Representative bioluminescence images of Rag2γc mice at day 7, showing limited tumor suppression by NEPN. These results demonstrate that NEPN significantly suppresses melanoma progression in immunocompetent B6 mice, but shows reduced efficacy in Rag2γc mice, suggesting that its anti-tumor function is partially dependent on an intact host immune system.

**Supplementary Figure 5.**
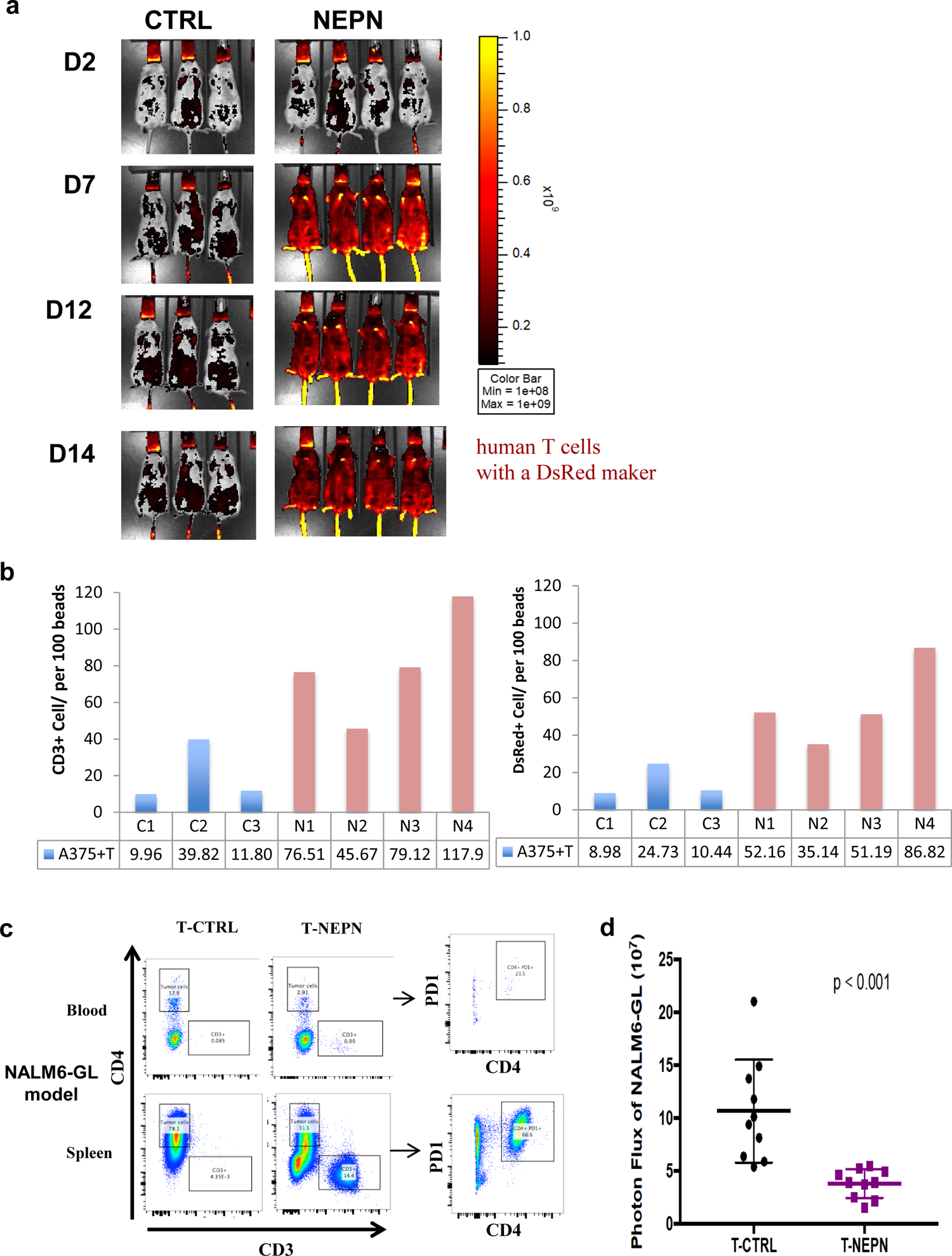
**a**, Bioluminescence imaging of A375 melanoma xenografts at serial time points following T cell transplantation with or without NEPN treatment. Proliferation signal of T-cells by measuring DsRed signal. **b,** Quantification of CD3⁺ and DsRed signal from individual mice of A375 melanoma xenografts at day 7 showing increased T cell proliferation in NEPN-treated groups compared to control. **c,** Flow cytometry analysis of peripheral T cell populations (CD3⁺, CD4⁺, PD1) from blood and spleen at day 18 from the mouse of Nalm6-GL xenografts, showing higher effector T cell persistence in NEPN-treated animals. **d,** Bioluminescent signals of leukemia NALM6-GL on day 7 in vivo xenograft model. P values calculated using the Mann-Whitney test.

**Supplementary Figure 6.**
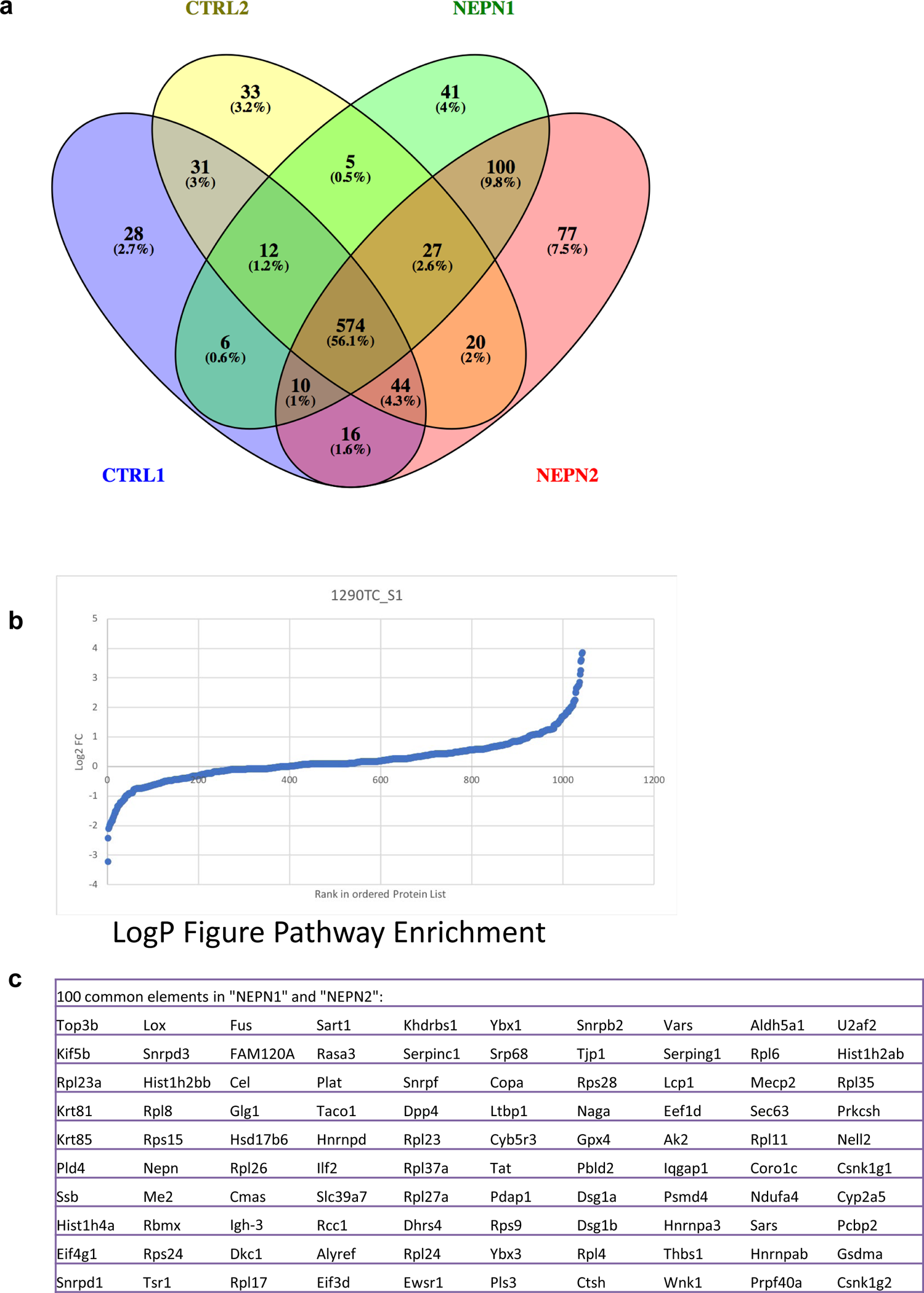
Identification of NEPN-interacting proteins by immunoprecipitation-mass spectrometry (IP-MS). **a,** Venn diagram showing overlap of proteins identified in independent NEPN pull-down experiments (NEPN1 and NEPN2) compared to control IPs (CTRL1 and CTRL2). A total of 574 proteins were shared across all four groups, while 100 proteins were specifically enriched in both NEPN1 and NEPN2 replicates, representing candidate NEPN-binding partners. **b,** Volcano/logP enrichment analysis of NEPN-associated proteins ranked by fold-change compared to control, illustrating significantly enriched interactors. **c,** List of the 100 common proteins specifically identified in both NEPN1 and NEPN2 datasets but absent in controls. These candidate interactors include proteins involved in RNA processing (Snrnp family, Rpl proteins), cytoskeletal remodeling (Tsr1, Krt family), metabolic regulation (Arg1, Aldh5a1, Gpx4), and signaling pathways (Hnrnp family, Rasa3, Serping1). Together, these data indicate that NEPN binds a distinct set of extracellular and intracellular regulatory proteins within the tumor microenvironment. This set includes LRBP, a candidate component of the ligand-receptor binding network, providing mechanistic support for the hypothesis that NEPN regulates HSP27- and ARG1-associated signaling through modulation of upstream cytokine or receptor pathways.

**Supplementary Figure 7.**
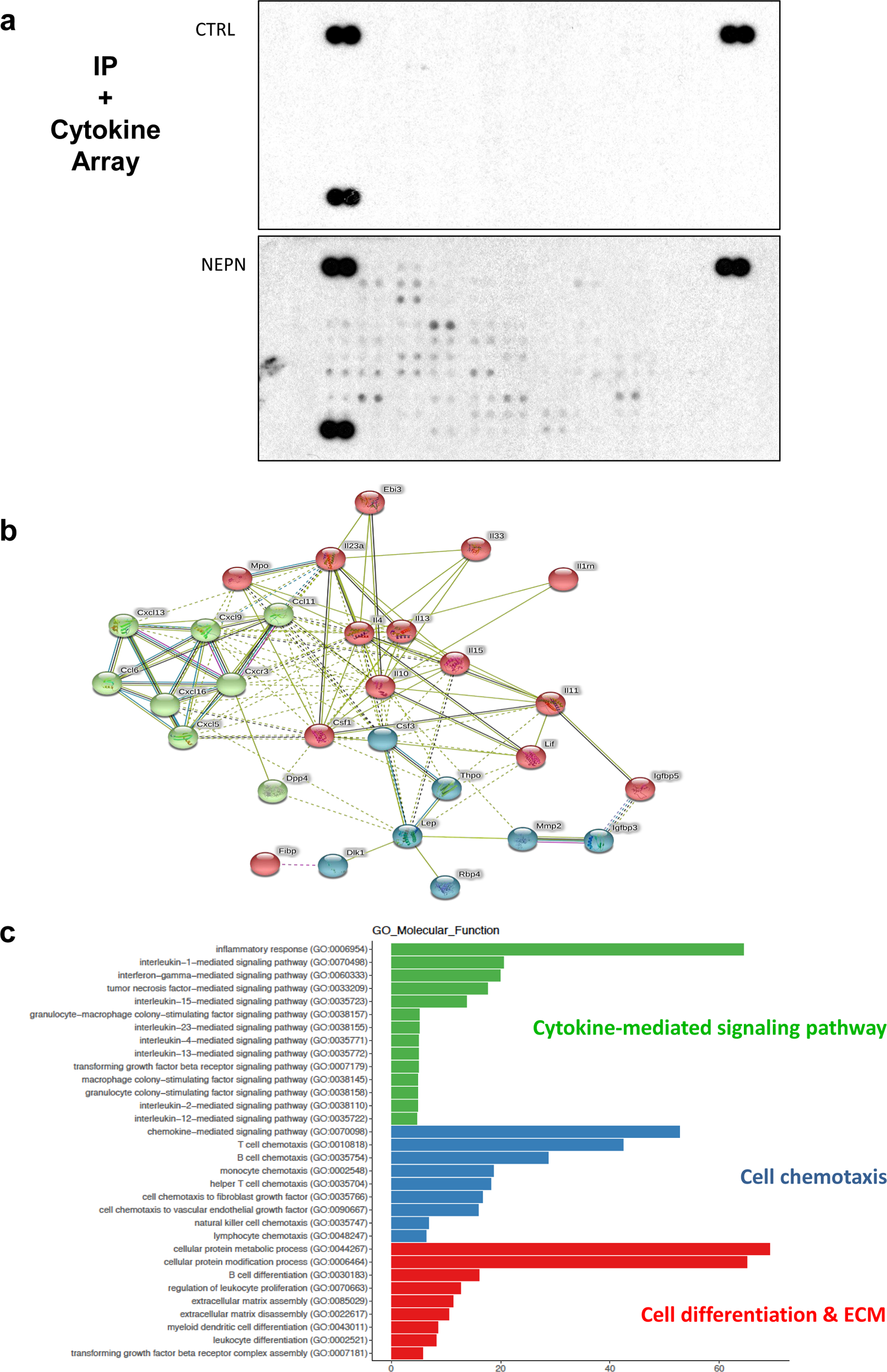
NEPN-interacting cytokines and pathway analysis. **a,** NEPN-interacting cytokines in the liver metastasis site were identified using immunoprecipitation followed by cytokine array (n = 4 independent experiments). **b,** Predicted cytokine network mediated by NEPN. **c,** NEPN-mediated signaling pathways were analyzed using pathway enrichment analysis, including cytokine-cytokine receptor interaction, chemokine signaling, T cell receptor signaling, immune response, and extracellular matrix organization. Among the identified cytokines, both IL-10 and TGF-β are known to cooperate in promoting regulatory T cell (Treg) induction and maintenance. This suggests that NEPN may modulate the immune environment not only by regulating effector immune cell chemotaxis but also by influencing Treg activity, a hypothesis to be tested in future studies.

## References

1 Slamon, D. J. & Cline, M. J. Expression of cellular oncogenes during embryonic and fetal development of the mouse. Proceedings of the National Academy of Sciences of the United States of America 81, 7141–7145 (1984).

2 Giuffrida, D. et al. Human embryonic stem cells secrete soluble factors that inhibit cancer cell growth. Cell proliferation 42, 788–798, doi:10.1111/j.1365-2184.2009.00640.x (2009).

3 Gootwine, E., Webb, C. G. & Sachs, L. Participation of myeloid leukaemic cells injected into embryos in haematopoietic differentiation in adult mice. Nature 299, 63–65 (1982).

4 Gerschenson, M., Graves, K., Carson, S. D., Wells, R. S. & Pierce, G. B. Regulation of melanoma by the embryonic skin. Proceedings of the National Academy of Sciences of the United States of America 83, 7307–7310 (1986).

5 Podesta, A. H., Mullins, J., Pierce, G. B. & Wells, R. S. The neurula stage mouse embryo in control of neuroblastoma. Proceedings of the National Academy of Sciences of the United States of America 81, 7608–7611 (1984).

6 Wells, R. S. & Miotto, K. A. Widespread inhibition of neuroblastoma cells in the 13- to 17-day-old mouse embryo. Cancer research 46, 1659–1662 (1986).

7 Webb, C. G., Gootwine, E. & Sachs, L. Developmental potential of myeloid leukemia cells injected into midgestation embryos. Developmental biology 101, 221–224 (1984).

8 Iozzo, R. V. & Schaefer, L. Proteoglycan form and function: A comprehensive nomenclature of proteoglycans. Matrix biology: journal of the International Society for Matrix Biology 42, 11–55, doi:10.1016/j.matbio.2015.02.003 (2015).

9 Frey, H., Schroeder, N., Manon-Jensen, T., Iozzo, R. V. & Schaefer, L. Biological interplay between proteoglycans and their innate immune receptors in inflammation. The FEBS journal 280, 2165–2179, doi:10.1111/febs.12145 (2013).

10 Iozzo, R. V. & Sanderson, R. D. Proteoglycans in cancer biology, tumour microenvironment and angiogenesis. J Cell Mol Med 15, 1013–1031, doi:10.1111/j.1582-4934.2010.01236.x (2011).

11 Schaefer, L. & Iozzo, R. V. Biological functions of the small leucine-rich proteoglycans: from genetics to signal transduction. The Journal of biological chemistry 283, 21305–21309, doi:10.1074/jbc.R800020200 (2008).

12 Mochida, Y. et al. Nephrocan, a novel member of the small leucine-rich repeat protein family, is an inhibitor of transforming growth factor-beta signaling. The Journal of biological chemistry 281, 36044–36051, doi:10.1074/jbc.M604787200 (2006).

13 Ghatak, S. et al. Roles of Proteoglycans and Glycosaminoglycans in Wound Healing and Fibrosis. International journal of cell biology 2015, 834893, doi:10.1155/2015/834893 (2015).

14 Tian, X. et al. High-molecular-mass hyaluronan mediates the cancer resistance of the naked mole rat. Nature 499, 346–349, doi:10.1038/nature12234 (2013).

15 Overwijk, W. W. & Restifo, N. P. B16 as a mouse model for human melanoma. Curr Protoc Immunol **Chapter** 20, Unit 20 21, doi:10.1002/0471142735.im2001s39 (2001).

16 Clark, E. A., Golub, T. R., Lander, E. S. & Hynes, R. O. Genomic analysis of metastasis reveals an essential role for RhoC. Nature 406, 532–535, doi:10.1038/35020106 (2000).

17 Einarsdottir, B. O. et al. Melanoma patient-derived xenografts accurately model the disease and develop fast enough to guide treatment decisions. Oncotarget 5, 9609–9618, doi:10.18632/oncotarget.2445 (2014).

18 Muller-Sieburg, C. The puzzling origin of lymphocytes. Blood 119, 5609–5610, doi:10.1182/blood-2012-04-420737 (2012).

19 Yoder, M. C. Inducing definitive hematopoiesis in a dish. Nature biotechnology 32, 539–541, doi:10.1038/nbt.2929 (2014).

20 Yoshimoto, M. et al. Autonomous murine T-cell progenitor production in the extra-embryonic yolk sac before HSC emergence. Blood 119, 5706–5714, doi:10.1182/blood-2011-12-397489 (2012).

21 Boiers, C. et al. Lymphomyeloid contribution of an immune-restricted progenitor emerging prior to definitive hematopoietic stem cells. Cell Stem Cell 13, 535–548, doi:10.1016/j.stem.2013.08.012 (2013).

22 Garrido, C. et al. Heat shock proteins 27 and 70: anti-apoptotic proteins with tumorigenic properties. Cell cycle 5, 2592–2601 (2006).

23 Musiani, D. et al. Heat-shock protein 27 (HSP27, HSPB1) is up-regulated by MET kinase inhibitors and confers resistance to MET-targeted therapy. FASEB journal: official publication of the Federation of American Societies for Experimental Biology 28, 4055–4067, doi:10.1096/fj.13-247924 (2014).

24 Bronte, V., Serafini, P., Mazzoni, A., Segal, D. M. & Zanovello, P. L-arginine metabolism in myeloid cells controls T-lymphocyte functions. Trends in immunology 24, 302–306 (2003).

25 Tham, M. et al. Melanoma-initiating cells exploit M2 macrophage TGFbeta and arginase pathway for survival and proliferation. Oncotarget 5, 12027–12042, doi:10.18632/oncotarget.2482 (2014).

26 Zhou, X., Spittau, B. & Krieglstein, K. TGFbeta signalling plays an important role in IL4-induced alternative activation of microglia. Journal of neuroinflammation 9, 210, doi:10.1186/1742-2094-9-210 (2012).

27 Pengal, R. et al. Inhibition of the protein kinase MK-2 protects podocytes from nephrotic syndrome-related injury. Am J Physiol Renal Physiol 301, F509–519, doi:10.1152/ajprenal.00661.2010 (2011).

28 Mizutani, H. et al. HSP27 modulates epithelial to mesenchymal transition of lung cancer cells in a Smad-independent manner. Oncol Lett 1, 1011–1016, doi:10.3892/ol.2010.190 (2010).

29 Missotten, G. S. et al. Heat shock protein expression in the eye and in uveal melanoma. Investigative ophthalmology & visual science 44, 3059–3065 (2003).

30 Neuzillet, C. et al. Targeting the TGFbeta pathway for cancer therapy. Pharmacol Ther 147, 22–31, doi:10.1016/j.pharmthera.2014.11.001 (2015).

31 Busse, A. & Keilholz, U. Role of TGF-beta in melanoma. Curr Pharm Biotechnol 12, 2165–2175 (2011).

32 Yang, L., Pang, Y. & Moses, H. L. TGF-beta and immune cells: an important regulatory axis in the tumor microenvironment and progression. Trends in immunology 31, 220–227, doi:10.1016/j.it.2010.04.002 (2010).

33 Zhu, J. et al. Comparative genomics search for losses of long-established genes on the human lineage. PLoS Comput Biol 3, e247, doi:10.1371/journal.pcbi.0030247 (2007).

34 Geiger, R. et al. L-Arginine Modulates T Cell Metabolism and Enhances Survival and Anti-tumor Activity. Cell 167, 829–842 e813, doi:10.1016/j.cell.2016.09.031 (2016).

35 Rubtsov, Y. P. & Rudensky, A. Y. TGFbeta signalling in control of T-cell-mediated self-reactivity. Nat Rev Immunol 7, 443–453, doi:10.1038/nri2095 (2007).

36 Saito, T. et al. Two FOXP3(+)CD4(+) T cell subpopulations distinctly control the prognosis of colorectal cancers. Nat Med 22, 679–684, doi:10.1038/nm.4086 (2016).

37 Flavell, R. A., Sanjabi, S., Wrzesinski, S. H. & Licona-Limon, P. The polarization of immune cells in the tumour environment by TGFbeta. Nat Rev Immunol 10, 554–567, doi:10.1038/nri2808 (2010).

38 Li, M. O. & Flavell, R. A. TGF-beta: a master of all T cell trades. Cell 134, 392–404, doi:10.1016/j.cell.2008.07.025 (2008).

39 Papapetrou, E. P. et al. Genomic safe harbors permit high beta-globin transgene expression in thalassemia induced pluripotent stem cells. Nature biotechnology 29, 73–78, doi:10.1038/nbt.1717 (2011).

40 Sadelain, M., Papapetrou, E. P. & Bushman, F. D. Safe harbours for the integration of new DNA in the human genome. Nature reviews. Cancer 12, 51–58, doi:10.1038/nrc3179 (2012).

41 Jiang, P. et al. Signatures of T cell dysfunction and exclusion predict cancer immunotherapy response. Nat Med, doi:10.1038/s41591-018-0136-1 (2018).

42 Suzuki, A., Leland, P., Joshi, B. H. & Puri, R. K. Targeting of IL-4 and IL-13 receptors for cancer therapy. Cytokine 75, 79–88, doi:10.1016/j.cyto.2015.05.026 (2015).

43 Teng, M. W., Darcy, P. K. & Smyth, M. J. Stable IL-10: a new therapeutic that promotes tumor immunity. Cancer cell 20, 691–693, doi:10.1016/j.ccr.2011.11.020 (2011).

44 Levings, M. K., Bacchetta, R., Schulz, U. & Roncarolo, M. G. The role of IL-10 and TGF-beta in the differentiation and effector function of T regulatory cells. Int Arch Allergy Immunol 129, 263–276, doi:10.1159/000067596 (2002).

45 Crowley, L. C., Christensen, M. E. & Waterhouse, N. J. Measuring Survival of Adherent Cells with the Colony-Forming Assay. Cold Spring Harbor protocols 2016, pdb.prot087171, doi:10.1101/pdb.prot087171 (2016).

46 Skamagki, M. et al. ZSCAN10 expression corrects the genomic instability of iPSCs from aged donors. Nat Cell Biol 19, 1037–1048, doi:10.1038/ncb3598 (2017).

47 Lee, Y. S. et al. Downregulation of NFAT2 promotes melanogenesis in B16 melanoma cells. Anatomy & cell biology 43, 303–309, doi:10.5115/acb.2010.43.4.303 (2010).

48 Riviere, I., Brose, K. & Mulligan, R. C. Effects of retroviral vector design on expression of human adenosine deaminase in murine bone marrow transplant recipients engrafted with genetically modified cells. Proceedings of the National Academy of Sciences of the United States of America 92, 6733–6737 (1995).

49 Gong, M. C. et al. Cancer patient T cells genetically targeted to prostate-specific membrane antigen specifically lyse prostate cancer cells and release cytokines in response to prostate-specific membrane antigen. Neoplasia (New York, N.Y.) 1, 123–127 (1999).

